# A shared manifold for scalable temporal representations in self-paced timing

**DOI:** 10.64898/2026.09.03.749079

**Authors:** Melissa Serrano, Manfredi Castelli, Yangfan Peng, Andrew Sharott, David Dupret

## Abstract

Adaptive behaviour often relies on tracking the passage of time, yet how distinct brain regions generate coherent temporal representations remains unclear. Using a self-paced interval timing task in mice, we show that heterogeneous single-neuron temporal firing profiles distributed across regions are organized within a shared ring manifold. Within this low-dimensional space, population activity evolves along a common trajectory across different intervals and encodes elapsed time in a relative reference frame. Different durations are not represented by separate neural states, but by modulation of traversal speed. These scalable dynamics arise from coordinated population-wide co-scaling of single-neuron activity and support trial-by-trial adjustments in timing behaviour. A cross-regional assembly of start neurons predicts, at interval onset, upcoming waiting duration and behavioural adjustments, linking initial population states to trajectory evolution. Together, these findings identify population traversal of a shared activity manifold as a mechanism for scalable temporal representation across brain regions.

## Introduction

Everyday behaviour requires generating actions at appropriate times. Whether playing an instrument or crossing a busy street, we rely on an internal estimate of elapsed time rather than merely reacting to external cues. This ability to estimate durations in the seconds to minutes range – termed interval timing – requires neural circuits to support a representation of elapsed time that provides a coherent temporal reference while remaining flexible across varying behavioural demands^1–3^. How such a reference emerges from neural population activity remains unclear.

A key challenge in addressing this question may lie in the distributed and heterogeneous nature of time representations. Neural activity correlated with elapsed time has been observed across numerous brain regions, including frontal and parietal cortices, striatum, thalamus, and hippocampus^4–10^. These signals are heterogeneous at the single-neuron level, encompassing ramping activity, sequential activation, and transient responses^4,6,8,10–15^. Because most studies have examined individual regions in isolation^5,7,8,16–18^, it remains unclear whether these temporal representations reflect distinct region-specific mechanisms or ubiquitous, coordinated population-level processes spanning multiple areas.

Recent work in motor and cognitive systems suggests that complex behaviours are supported by low-dimensional population dynamics embedded within high-dimensional neural activity^19–25^. In these frameworks, behaviour emerges from structured trajectories through low-dimensional state spaces rather than static representations in individual neurons^26–28^. Whether timing behaviour is implemented through similar dynamics across circuits, and how such dynamics are coordinated, remains unknown.

Here we developed a self-paced interval timing task in which mice generated reaching movements at specific target delays, enabling precise measurement of timing behaviour as a readout of internal time estimates. Using simultaneous high-density recordings across multiple regions, we examined how elapsed time is represented in brain-distributed neurons on individual trials. Applying non-linear dimensionality reduction to this population activity, we derived a low-dimensional embedding to test whether heterogeneous timing-related responses are organized within a shared manifold. We then examined whether different interval durations are encoded by changes in neural states or by scalable modulation of population trajectories, and how coordinated activity predicts trial-by-trial behavioural adjustments. Together, these results identify population co-scaling of distributed temporal representations within a shared ring manifold as a unifying mechanism for the flexible internal estimation of elapsed time.

## Results

### Learning to time behaviour to a target interval

We developed a self-paced interval timing task in which head-fixed mice learned to initiate reaching movements after specified target delays to obtain water rewards (Fig. 1a). Each session was assigned a distinct target delay in the seconds range (for example, 3 seconds). Mice initiated trials by touching a handbar, which they were required to hold until the target delay elapsed before releasing and reaching toward a reward spout (Fig. 1b). Hold durations were centred near the session target interval (Figs. 1c and S1a-b), with a slight bias above the target (Fig. 1d; mean normalised lag from target: 1.25; 95% CI: 1.16, 1.34). Across training, hold duration distributions became progressively steeper around each target delay (Fig. S1c,-d), showing that mice increased the proportion of near-target trials within each session.

**Fig. 1:**
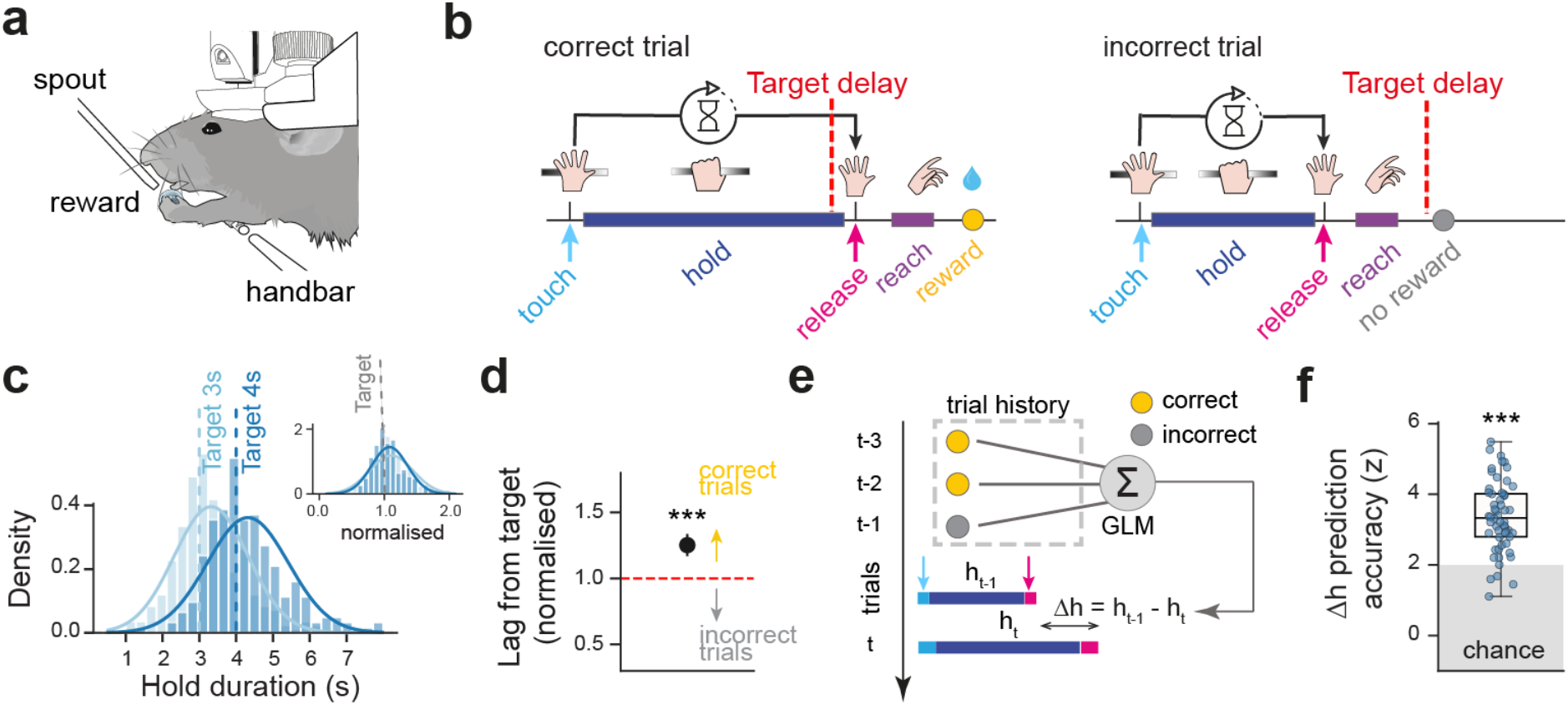
Behavioural performance and trial history-dependent timing in a self-paced interval task. **(a)** Experimental setup. Head-fixed mice initiated trials by contacting a handbar and obtained a sucrose reward by releasing and reaching after a target interval. **(b)** Task structure. Trials began at handbar touch. Release after the session-specific target delay triggered reward delivery (correct trial, left); premature release yielded no reward (incorrect trial, right). **(c)** Hold-duration distributions from two sessions of the same mouse. Dashed vertical lines indicate target delays. Inset, distributions normalized to each session target delay, showing overlap between sessions with distinct target delays. **(d)** Mean target-normalized hold duration per session. Mean duration tightly exceeded the target (mean target-normalised duration > 1, p < 10⁻⁵, one-tailed bootstrap test). Red dashed line indicates the target delay. **(e, f)** Trial-history influence on timing behaviour adjustments. **e**, Schematic of regression models using outcomes of the three preceding trials (t-1, t-2, t-3) to predict trial-to-trial changes in hold duration (Δh). **f**, Session-wise prediction accuracy, normalized to shuffled controls (mean (95% CI) accuracy: 3.36 (3.10–3.61) z; accuracy > 1.95, p < 10⁻⁵, one-tailed bootstrap test). See also Fig. S1e,f. *** p < 0.001. Data in d–f: n = 56 sessions from 9 mice.

We next examined whether timing behaviour was adjusted according to recent trial history rather than varying randomly across trials. Linear regression models predicting trial-by-trial changes in hold duration from reward history (Fig. 1e) outperformed shuffled controls (Fig. 1f). Adjustments were best predicted by the immediately preceding trial, with weaker contributions from earlier trials (Fig. S1e). Incorrect (unrewarded) trials were followed by longer holds, whereas correct (rewarded) trials were followed by shorter holds (Fig. S1e-f). Thus, mice learned to time behaviour to a target interval and flexibly updated their elapsed time estimate on a trial-by-trial basis to guide behaviour, providing a foundation for studying its associated neural representation.

### Population dynamics during interval timing are distributed and heterogeneous

To characterise neural population activity during interval timing, we performed high-density recordings using multiple Neuropixels probes simultaneously across prefrontal cortex, motor cortex, dorsal striatum, nucleus accumbens, parietal cortex, hippocampus, and posterior thalamus in trained mice [Fig. 2a and Table S1; total 8,642 neurons from 31 sessions, 6 mice; mean (IQR) number of neurons per session: 278.8 (233.5–318.0)]. These regions were selected to sample key regions individually investigated in the context of time perception. Previous work has consistently implicated prefrontal, motor and parietal cortices in temporal estimation and the control of timed actions^3,6,15,16,29^ . The role of the striatum and broader basal ganglia in interval timing is supported by lesion, pharmacological and electrophysiological studies in animals, as well as human neuroimaging^2,3,7,30–32^. The presence of sequential time cells and ramping cells suggests that the hippocampus might contribute to the representation of elapsed time^8,33,34^. Finally, higher-order thalamic nuclei form extensive reciprocal connections with cortical, hippocampal and striatal circuits, supporting communication and the coordination of distributed computations that may underlie temporal behaviour^9,35–37^. Activity from the same neurons was also recorded during off-task control periods, when mice could hold the handbar but were not engaged in interval timing (Fig. S2a).

**Fig. 2:**
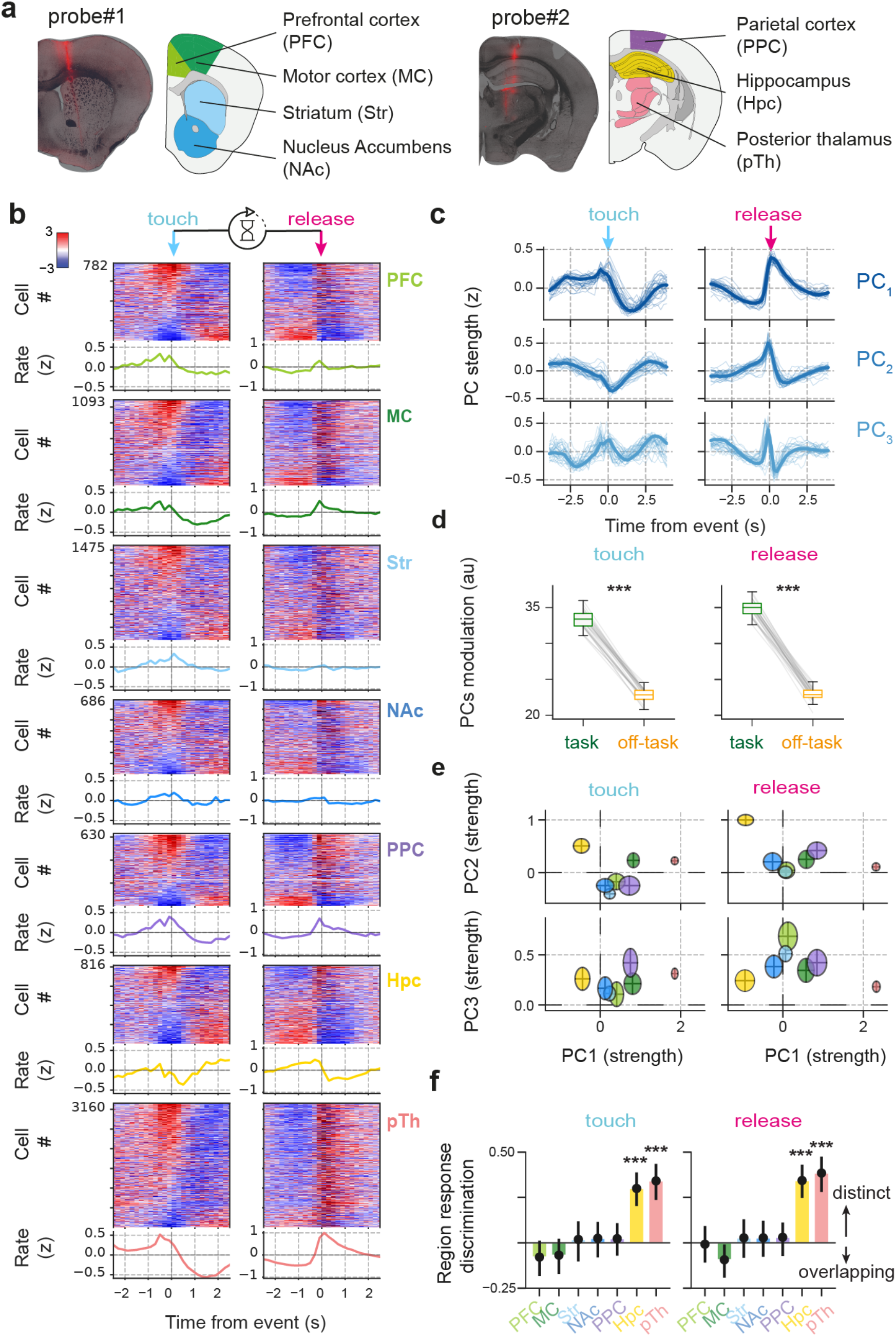
Distributed and heterogeneous population responses across brain regions during interval timing. **(a)** Neuropixels probe placements showing recorded regions and representative probe trajectories from an example mouse. **(b)** Triggered average firing responses across regions, aligned to handbar touch (left) and handbar release (right). For each region, top: z-scored firing rates of individual neurons sorted by activity at handbar touch; bottom: mean population response. PFC, prefrontal cortex; MC, motor cortex; Str, striatum; NAc, nucleus accumbens; PPC, posterior parietal cortex; Hpc, hippocampus; pTH, posterior thalamus. **(c)** First three principal components (PCs) of responses in **b**, computed across regions. Rows: PCs; columns: alignment to handbar touch (left) and release (right). Thin lines: individual sessions; thick lines: full dataset. **(d)** Session-mean PCs modulation strength during task versus off-task control epochs for handbar touch and release (handbar touch, p < 10⁻⁵; handbar release, p < 10⁻⁵; paired bootstrap tests). **(e)** Projection of neurons from each region into the PC space shown in **c**. Ellipses denote 95% confidence intervals. **(f)** Linear discriminant analysis of PC projections for region classification. Cross-validated scores were normalised to shuffled controls. Hpc and pTH exceeded chance (p < 10⁻^3^, one-tailed bootstrap test; Bonferroni corrected); other regions did not (all Ps > 0.15). Error bars, 95% CI across 1,000 permutations with equal sampling of neurons per region. *** p < 0.001. Data in b–f: n = 31 sessions from 6 mice.

We focused on the hold interval, defined as the period between handbar touch and handbar release (Fig. 1b), during which mice were required to hold the handbar and thus did not move. Trial-averaged firing rates aligned to either task event exhibited temporal modulation beyond transient event-locked responses (Fig. 2b). To characterise the dominant patterns of these responses across the recorded neuronal population, we applied principal component analysis (PCA) to the full time-course of these hold-duration responses. These PCs were consistent across sessions and mice and included monotonic ramping as well as non-monotonic event-locked responses (Figs. 2c and S2b). These patterns persisted after excluding trials that immediately followed reward presentation (Fig. S2c) and were weaker during duration-matched off-task windows (Fig. 2d).

All regions exhibited such population-level modulation (Fig. S2d). To assess their individual contributions to the global patterns, we projected region-specific responses into the population-wide PC activity space. Single-region responses occupied distinct yet partially overlapping positions within this common space (Fig. 2e), with hippocampus and thalamus showing the greatest separation (Fig. 2f). This cross-regional heterogeneity was stronger during task than off-task periods (Fig. S2e,f). These results show that population dynamics during interval timing were widely distributed across brain regions but nonetheless exhibited consistent, task-dependent heterogeneity.

### Population activity across brain regions carries information about elapsed time

To examine whether population responses carried task-relevant temporal information, we used non-linear regression models to decode, from neural activity on each hold interval, the elapsed time since handbar touch (Fig. 3a). We trained models to predict either absolute time (the physical time elapsed, for example 1 s from trial start) or relative time (the fraction of the interval elapsed, for example 10% of total trial duration). Both models performed above surrogate models trained on shuffled time; relative time, however, showed higher accuracy (Fig. 3b,c). A similar higher decoding performance for relative time was observed at the single-neuron level (Fig. S3a,b). Decoding remained at chance level during off-task windows for both elapsed time models (Fig. 3d,e). Subsequent analyses therefore focused on relative time.

**Fig. 3:**
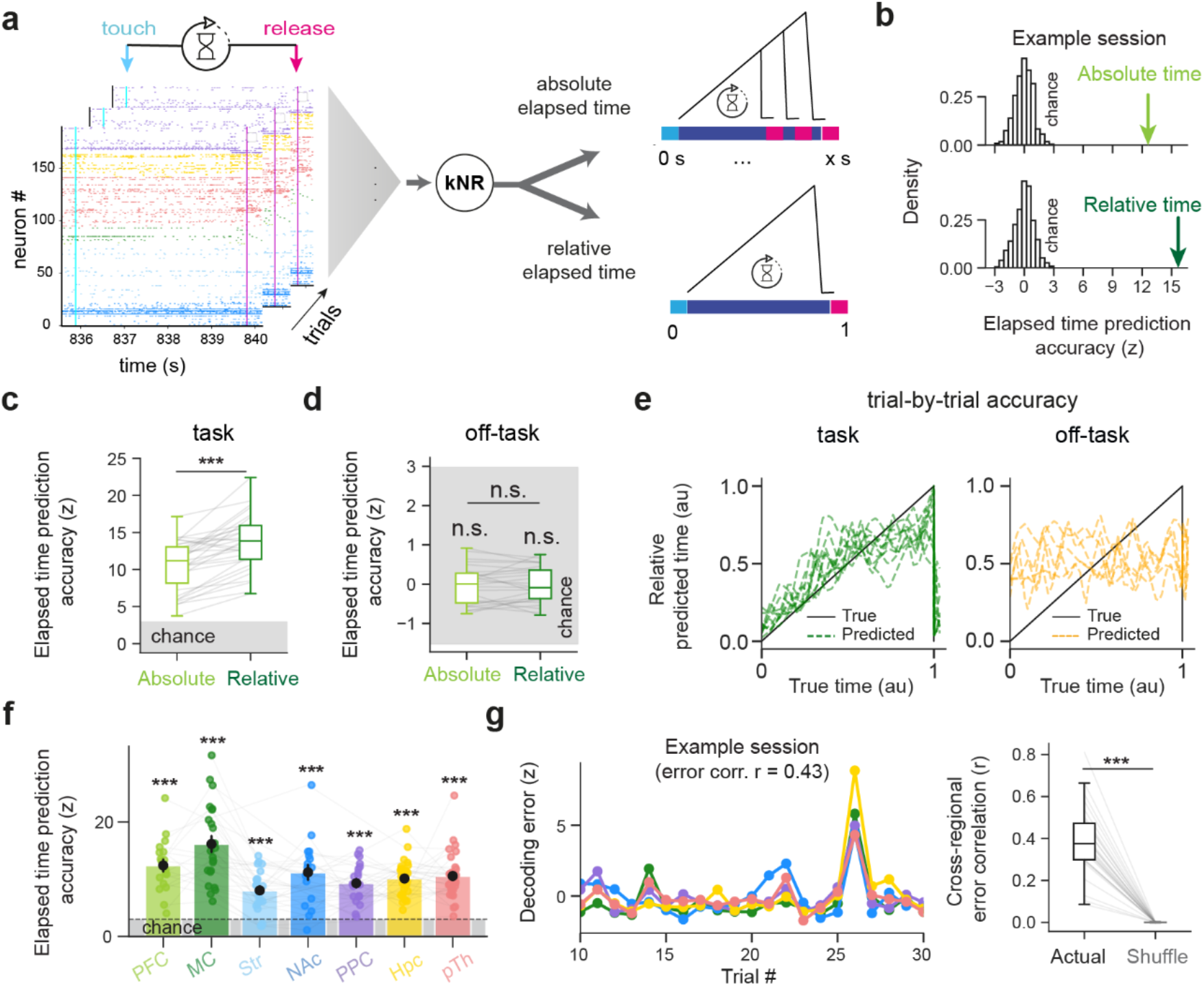
Population activity encodes elapsed time across brain regions. **(a)** Decoding of elapsed time from neural activity. Trial-by-trial activity from handbar touch to 200 ms before release (to exclude activity immediately preceding movement) was used to train k-nearest neighbours regression models (kNR) to predict absolute or relative elapsed time. **(b)** Chance-normalized decoding accuracy from an example session for absolute (top) and relative (bottom) elapsed time. Accuracies were z-scored relative to shuffled time. **(c, d)** Session chance-normalized decoding accuracy during task (c) and off-task control windows (d). During task, both decoding models exceeded change (accuracy > 1.96 z, all Ps < 10^-5^; one tailed bootstrap test), and relative time decoding exceeded absolute time (p < 10⁻⁵; paired bootstrap test). During off-task windows, neither decoding exceeded chance (accuracy > 1.96 z, all Ps = 1; one tailed bootstrap test) and there was no difference between absolute and relative time decoding (p = 0.78; paired bootstrap test). Shaded area indicates chance level. **(e)** Example trial-by-trial prediction of relative elapsed time during task and off-task intervals. **(f)** Region-wise decoding chance-normalised accuracy for relative elapsed time. All regions exceeded chance (normalised accuracy > 1.96 z, p < 10⁻^5^; one-tailed bootstrap tests; Bonferroni corrected). Accuracy differed across regions (χ²(6) = 56.6, p = 2.2 × 10⁻^10^; likelihood ratio test from a linear mixed-effects model with region as fixed effect and session as random intercept), with highest values in MC. Shaded area indicates chance level. **(g)** Cross-regional co-fluctuation of relative-time decoding errors. Left, example trial-by-trial decoding errors across (color-coded) regions. Right, session-wise error correlations exceeded shuffled controls (p < 10⁻⁵; one-tailed paired bootstrap test). *** p < 0.001. Data in c,d,f,g: task, n = 31 sessions from 6 mice; off-task, n = 26 sessions from 5 mice.

To test whether relative elapsed time could be decoded from each region’s activity, we trained independent region-specific models (controlling for differences in neuron count across regions and sessions). Decoding accuracy exceeded chance in all regions (Fig. 3f), with prefrontal and motor cortices showing higher performance ^5,29,38^. Decoding errors were positively correlated across regions and this cross-regional correlation exceeded that observed in surrogate data, where regions decoded elapsed time independently of one another (Figs. 3g and S3c,d). Thus, elapsed time was preferentially encoded in a relative time frame across all regions during self-paced timing and exhibited coordinated decoding fluctuations.

### Organisation of single-neuron temporal responses within a shared manifold

During the hold interval, single neurons exhibited heterogeneous responses that could be summarised by six temporal activity profiles (Fig. 4a). Start neurons showed a sharp increase in firing at trial initiation, coinciding with handbar touch (i.e., interval-onset cells). Two additional profiles featured monotonic changes throughout the hold period: ramp-up neurons gradually increased their activity, whereas ramp-down neurons decreased firing. Middle neurons displayed peak firing at a consistent intermediate time within the hold interval across trials (i.e., mid-interval cells). End neurons were transiently activated at handbar release (i.e., interval-offset cells). Bracket neurons displayed two bursts of activity near handbar touch and handbar release.

**Fig. 4:**
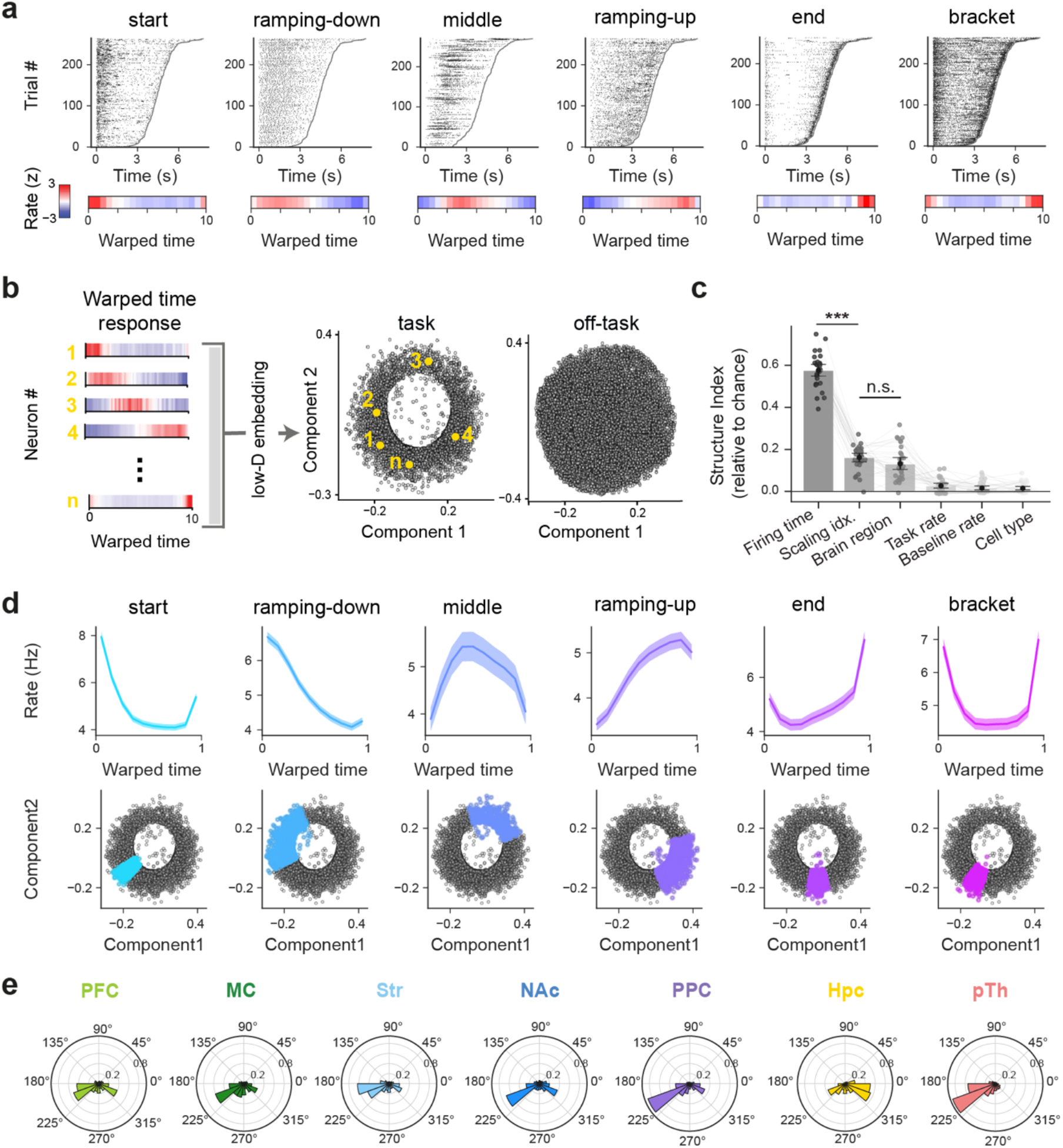
Single-neuron temporal responses are organized within a low-dimensional ring manifold. **(a)** Example neurons exhibiting start, ramp-down, middle, ramp-up, end, and bracket firing response profiles. Top, spike rasters across trials sorted by duration (border line, trial end). Bottom, z-scored warped-time responses. Each column represents one neuron. **(b)** Low-dimensional manifold of warped-time responses. Left, dimensionality reduction schematic. Warped-time responses of all neurons were used as input to Isomap to obtain a low-dimensional manifold. Right, resulting Isomap manifold showing ring geometry during task but not during off-task control epochs. Each dot represents one neuron. Note that during task performance, neurons 1–5 (numbers in yellow) occupied distinct angular positions. **(c)** Structure index (SI) of manifold organisation for candidate latent variables. SI values were normalised to shuffled controls. Preferred peak firing time accounted for the largest contribution, followed by scaling index and brain region (difference across latent variables: χ²(5) = 499.9, p = 8 × 10⁻⁶; likelihood ratio test from a linear mixed-effects model with latent as a fixed effect and session as a random intercept). Each dot represents one session. Error bars, 95% CI. **(d)** Mean warped-time responses by response profile (top) and corresponding positions in the manifold (bottom). Angular range (response profile classifier accuracy > 95%): start 199-226°, ramp-down 107-204°, middle 11-110°, ramp-up 15-285°, end 249-288°, bracket 221-255°. Shaded areas, 95% CI. **(e)** Angular distribution of neurons in the ring manifold. Neurons from all regions were distributed across angular sectors, with region-specific biases. *** p < 0.001. Data in c: n = 31 sessions from 6 mice; b,d,e: n = 8,642 neurons, full dataset.

Applying non-linear dimensionality reduction to time-warped activity profiles revealed a ring-like population manifold, with neurons arranged along a continuum (Fig. 4a,b and S4a). This geometry was absent in off-task population activity (Figs. 4b and S4a-b). Moreover, during task performance, neurons retained consistent angular positions on manifolds constructed and evaluated across distinct sets of trials; this was not the case for off-task periods (Fig. S4c,d).

We next determined which latent variables best explained the geometry of the ring by computing the Structure Index^39^. Within-trial peak firing time accounted for most of the variance in the manifold organization, followed by the consistency of firing profiles across trials of different durations (quantified using the scaling index^26^), and then by region identity (Fig. 4c). Mean firing rate (during task performance and off-task baseline periods) and putative cell type (broad-spike principal neurons versus narrow-spike interneurons) contributed minimally (Fig. 4c). Peak firing time was mapped onto the angle within the ring such that neurons with similar angular positions exhibited similar temporal firing profiles (Fig. 4d), with all six response profiles (Fig 4a) distributed along this continuum. Elapsed-time decoding accuracy (Fig. 3) varied with angular position: ramping neurons showed the highest decoding performance, whereas middle neurons showed the lowest (Fig. S4e). Radial position within the manifold instead reflected the cross-trial response consistency of single-neurons (Fig. S4f). Neurons from all regions occupied sectors of the manifold corresponding to all six temporal profiles, but with region-specific biases in angular distribution (Figs. 4e and S4g). For example, start neurons were more prevalent in parietal cortex and posterior thalamus, ramping-up neurons in hippocampus, and end as well as bracket neurons in motor cortex. These results show that, while hold-duration related activity was distributed and heterogeneous, single-neuron response profiles are captured within a low-dimensional manifold shared across neurons.

### Population activity traverses the ring manifold as a scalable trajectory

Given that the angular position on the manifold reflected when each neuron preferentially fires within the trial, we next investigated whether, on a trial-by-trial basis, the manifold captured the progression of instantaneous population activity from hold duration start to end. In line with this, sorting neurons by angular position within the manifold revealed sequential activation across the interval on single trials (Fig. 5a). We thus projected each instantaneous population activity vector onto the ring manifold (Fig. S5a), yielding angular trajectories that evolved continuously over the full trial duration (Figs. 5b and S5b). A non-linear classifier trained to predict elapsed time from these trial-wise trajectories performed above chance (Fig. 5c). The classifier’s decision boundaries formed a fan-like partitioning of the state space aligned to angular progression (Fig. 5d), indicating that population activity orderly traversed the full ring consistently across trials. Importantly, this suggested that trials with different hold durations followed the same trajectory, differing only in traversal speed, with shorter trials progressing faster and longer trials more slowly (Figs. 5e-f and S5c). Therefore, population activity progressed along a common trajectory on the manifold in each trial, with the hold duration represented by traversal speed.

**Fig. 5:**
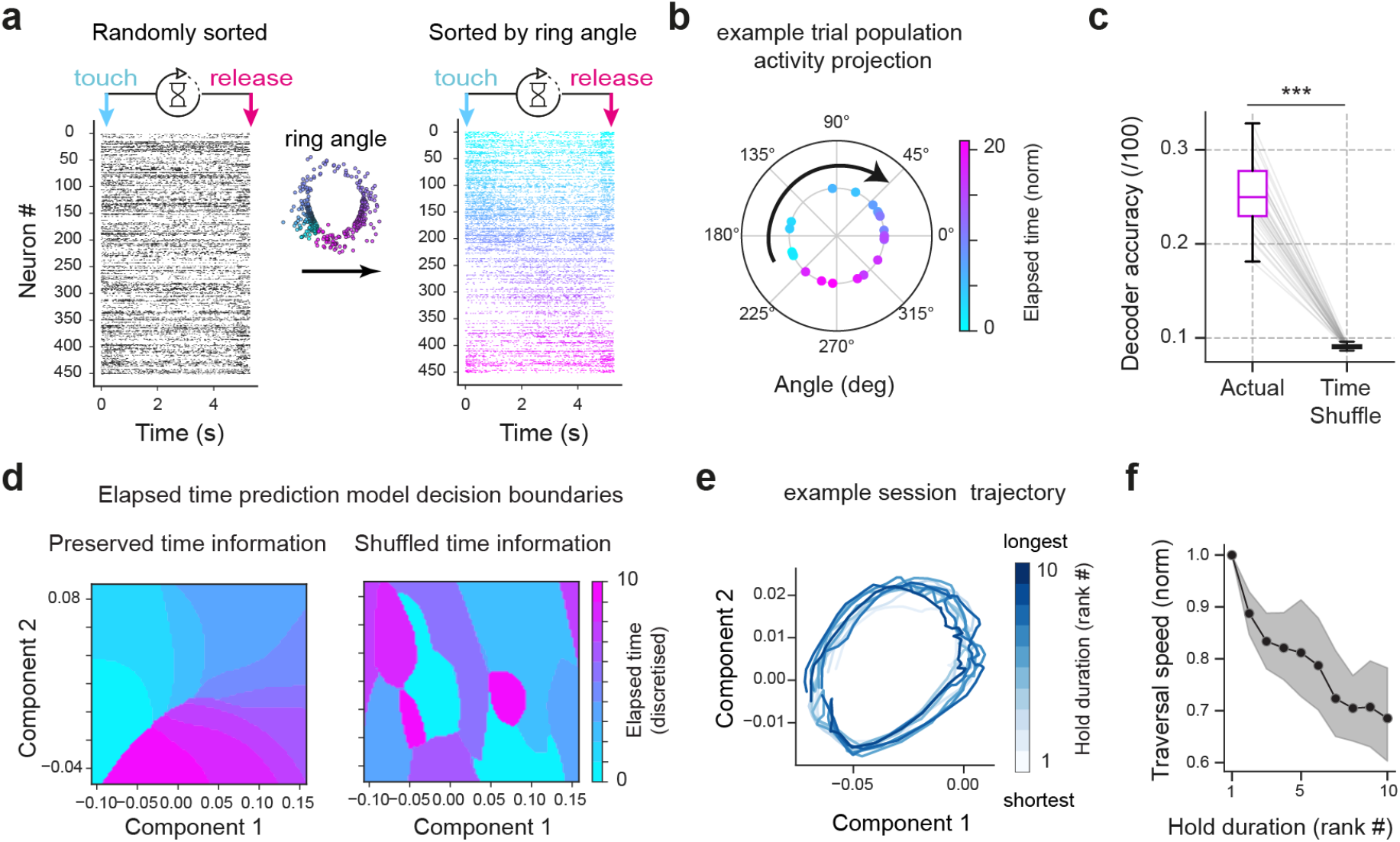
Population activity forms a low-dimensional temporal trajectory. **(a)** Trial-by-trial spike raster from one session between handbar touch and release. Recorded neurons (∼450; left) were sorted by their angular position in the ring manifold (right), relative to start neurons. **(b)** Projection of instantaneous population activity vectors onto angular coordinates of the manifold, yielding single-trial angular trajectories. Each trial unfolds along the ring (see also Fig. S5c). **(c, d)** Decoding of relative elapsed time from single-trial angular trajectories, using a support vector machine. **c**, Decoding accuracy exceeded surrogate models generated by shuffling elapsed time labels (p < 10⁻⁵; one-tailed paired bootstrap test). **d**, Example decision boundaries from one session using true labels (left) and shuffled labels (right). The model trained on true data exhibits a fan-like circular structure, which is absent in the shuffled control. **(e)** Population angular trajectories, averaged across ten bins of increasing trial duration from one session. Each color-coded trace represents the mean population activity projected in low-dimensional space from handbar touch to 0.5 s after release. Trajectories overlap despite differing durations, consistent with temporal scaling. **(f)** Normalized manifold traversal speed across trial duration ranks. Trials were binned by duration within session, and average manifold trajectories were computed per bin. *Shaded area*, 95% CI. Speed decreased with duration rank (Spearman r = −0.78, 95% CI [−0.93, - 0.59]; correlation different from zero: p < 10^-5^; bootstrap test). *** p < 0.001. Data in c-f: n = 31 sessions from 6 mice.

### Cross-regional population co-scaling during interval timing

Given that neural activity follows consistent trajectories across trials of different durations (Figs. 5e and S5c), we hypothesized that this consistency reflects coordinated temporal scaling of the diverse single-neuron firing profiles that jointly represent each trial. Under this framework, neurons would systematically change their activity patterns in proportion to the interval, preserving the trajectory geometry at the population level. We refer to such a hypothetical process as “population co-scaling”. Consistent with this idea^26,34,40^, we found that individual neurons exhibited temporal scaling across trials: start neurons modulated peak firing magnitude, ramping neurons adjusted response slope, and middle, end, and bracket neurons shifted peak response times (Fig. 6a). However, such single-neuron scaling does not necessarily imply coordinated adjustments across the population (co-scaling). Instead, similar population-level structure could arise from independent scaling in different subsets of neurons across trials (independent scaling). To distinguish between these two possibilities, we tested whether trial-by-trial adjustments reflected coordinated changes across the population rather than independent scaling of individual neurons (Fig. 6B). To do so, we used regression models that predicted each neuron’s activity changes from the rest of the population. Models preserving population co-scaling outperformed independent-scaling surrogates (Fig. 6c). More neurons exhibited co-scaling than expected by chance (Fig. S6a). Analysing the regression coefficients of these models revealed that population co-scaling relationships were not confined within anatomical regions: significant cross-regional predictive interactions were observed (Figs. 6d,e). These findings support a cross-regional co-scaling mechanism underlying consistent population temporal trajectories.

**Fig. 6:**
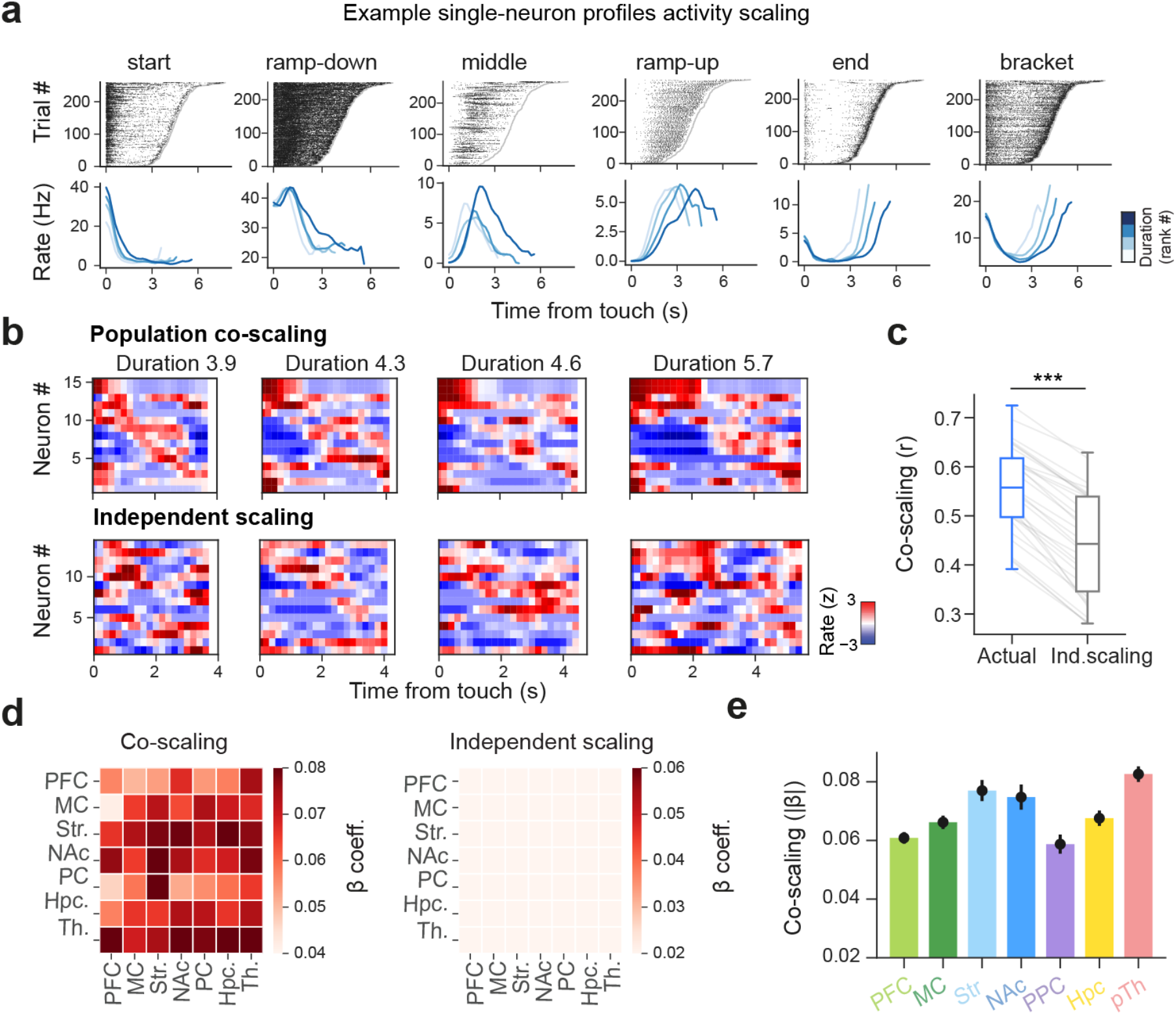
Cross-regional co-scaling of neuronal activity during interval timing. **(a)** Example single-neurons exhibiting start, ramp-down, middle, ramp-up, end, and bracket response profiles. Top, spike rasters across trials sorted by hold duration (vertical border, trial end). Bottom, mean spike counts aligned to handbar touch for quartiles of hold duration (light to dark blue). Each column represents to one neuron. **(b)** Firing responses (z-scored) from 15 example neurons across 4 trials of increasing duration (one trial per column). Top, neurons with coordinated scaling across durations (population co-scaling); bottom, neurons from the same trials but without consistent scaling. **(c–e)** Co-scaling analysis. **c**, Session-mean prediction accuracy for single-neuron activity from the rest of the population (leave-one-out), computed trial-by-trial. Actual data were compared to a surrogate distribution preserving individual neuron scaling statistics while shuffling co-scaling across neurons (independent scaling). Prediction magnitude exceeded surrogate control (p < 10⁻^5^; paired bootstrap test). **d**, Session-wise mean regression coefficients of scaling models grouped by region for significant co-scaling neurons. Rows, predicted neuron; columns, contributing regions. Left, actual coefficients; right, independent-scaling surrogate. **e**, Region-wise mean co-scaling strength derived from d. Information differed across regions, with the highest values in pTH and lower values in Hpc and several cortical regions (χ²(6) = 45.90, p = 3.10 × 10⁻⁸; likelihood ratio test from a linear mixed-effects model with region as a fixed effect and session as a random intercept). Error bars, 95% CI across neurons. *** p < 0.001. Data in c: n = 31 sessions from 6 mice. Data in d,e: n = 6,065 neurons.

### A cross-regional assembly of start-neurons predicts trial-by-trial timing behaviour

If population traversal speed varies across trials and is coordinated across regions, what determines this modulation? Single-neuron activity showed modulation associated with hold duration (Fig. 6a). We thus examined how ensemble activity of neurons grouped by temporal profiles related to the timing of reaching behaviour. Regression models were trained to predict total hold duration on individual trials from neuronal ensemble activity aligned to bar touch (Figs. 7a and S6b). The average decoding performance from these ensembles rose above chance shortly after handbar touch, within the first few hundred milliseconds, to then peak later in the trial (Fig. 7b). Importantly, information about total hold duration was already carried by the start-neuron ensemble at trial onset, and this was not the case for the other neuron ensembles (Fig. 7c,d and S6b).

**Fig. 7:**
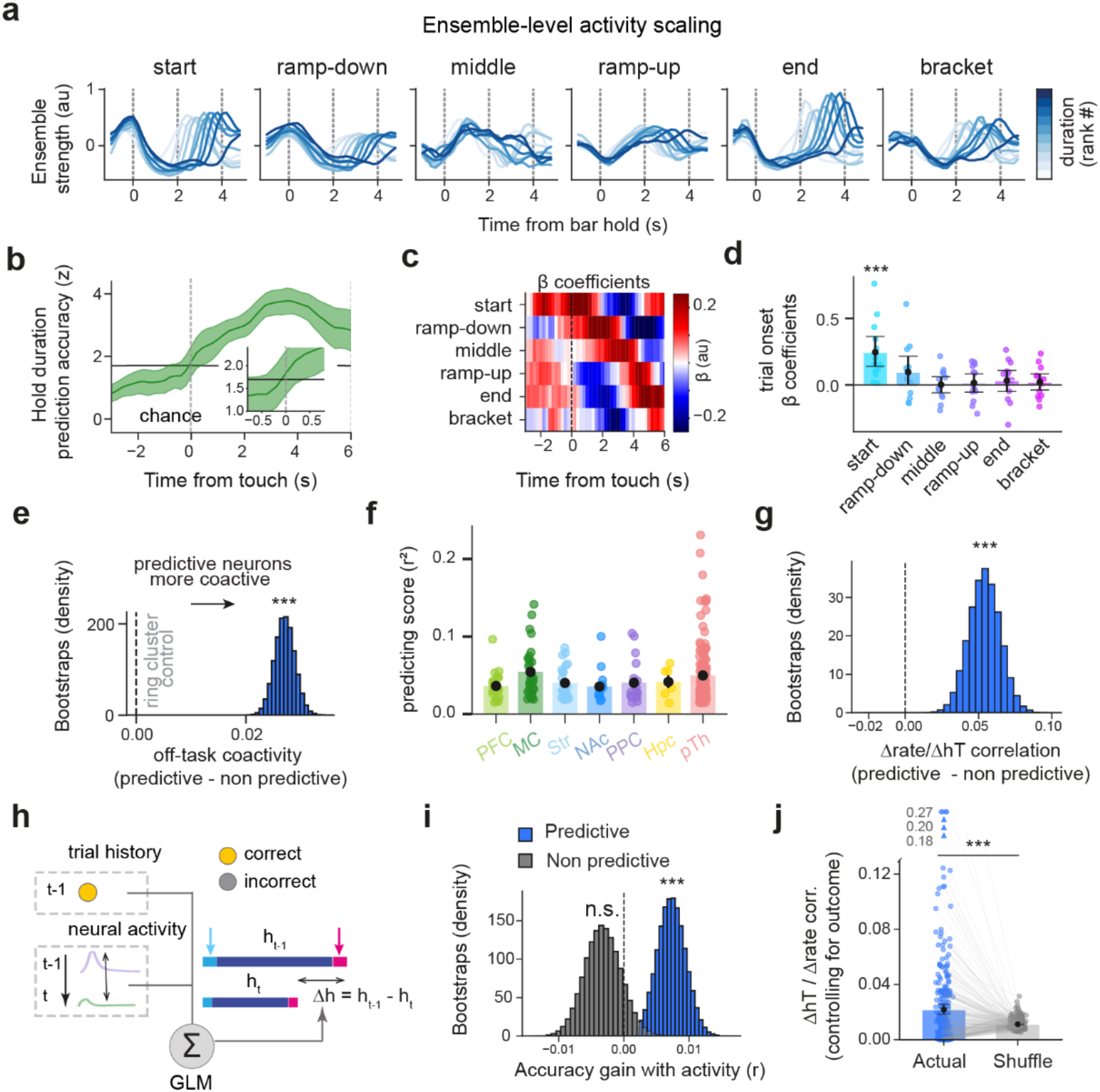
Distributed start-neuron activity predicts trial-by-trial interval timing and behavioural adjustments. **(a)** Ensemble-level firing responses aligned to handbar touch for trials of different durations from one session. Columns show mean activity profiles for ensembles of start, ramp-down, middle, ramp-up, end, and bracket neurons, across ten ranks of increasing trial duration. **(b–d)** Prediction of trial-by-trial duration from ensemble activity aligned to handbar touch (as in a), using linear regression. **b**, Mean decoding accuracy over time relative to handbar touch. Accuracy exceeded chance (estimated from surrogate models) after movement onset. Shaded area, 95% CI. **c**, Time-resolved regression coefficients from the model in b. The start-neuron ensemble showed elevated coefficients within hundreds of milliseconds after trial onset. **d,** Session-wise regression coefficients for each ensemble in an early decoding window after handbar touch (0–200 ms). Error bars, 95% bootstrap CI. Only the start ensemble showed beta coefficients significantly above zero (mean beta different from zero: start, p < 10^-5^; other ensembles, all Ps > 0.34; bootstrap tests, Bonferroni corrected). **(e)** Bootstrapped mean difference in off-task coactivity between hold-duration-predictive versus non-predictive start neurons (p < 10^-5^; bootstrap test; n = 264 predictive neurons, 257 non-predictive neurons from 5 mice). **(f)** Single-neuron prediction scores for hold duration among start neurons across regions. Information was distributed across all regions, with highest information observed in MC and pTH (χ²(6) = 70.68, p = 2.97 × 10⁻¹³; likelihood ratio test from a linear mixed-effects model with region as a fixed effect). **(g)** Cross-trial correlation of activity for predictive is higher than that of non-predictive start neurons (p < 10^-5^, bootstrap test; n = 297 predictive neurons, 290 non-predictive neurons from 6 mice). **(h–j)** Behavioural model incorporating neural activity. **h**, Linear regression model predicting trial-to-trial changes in hold duration (Δh) from both previous trial outcome and the change in firing rate of hold-duration-predictive (versus non-predictive) start neurons. **i**, Change in model performance relative to outcome-only model, compared to shuffled controls. Adding activity from predictive start neurons improved performance (p = 2 x 10^-4^; paired bootstrap tests; n = 28 sessions from 6 mice), whereas non-predictive neurons did not (p = 0.80; paired bootstrap test). **j**, Predictive power (R²) between changes in firing rate of hold-duration– predictive start neurons and changes in hold duration across consecutive trials with identical outcomes (e.g., two consecutive correct or incorrect trials). Each dot represents one neuron. Changes in firing rate predicted duration differences beyond chance for same-outcome trials, estimated from surrogate data generated by circularly shuffling trial duration (p < 10^-5^; paired bootstrap test; n = 297 predictive neurons from 6 mice). *** p < 0.001. Data in d,e: n = 6,065 neurons; in b–d: n = 13 sessions from 6 mice.

Given this early predictive signal at trial onset, we focused on start neurons and tested whether they exhibited coordinated activity. To avoid confounds from task covariates, pairwise millisecond-timescale coactivity among distributed start neurons was assessed during off-task periods. Hold-duration predictive start neurons showed stronger coactivity than non-predictive start neurons (Fig. 7e). The expression of this assembly of coactive start neurons was independent of task-related firing rate differences (Fig. S6c). Start neurons from all recorded regions contained information at trial onset about the total hold duration of the current trial, with highest predictive strength observed in motor cortex, posterior parietal cortex, and thalamus (Figs. 7f and S6d).

We observed that total hold duration was adjusted according to previous trial’s outcome (Fig. 1e,f). We thus tested whether hold-duration predictive start-neuron assembly activity also contributed to such trial-by-trial behavioural adjustment. Activity among these start neurons was more strongly correlated with trial-by-trial adjustments in hold time than activity among non-predictive start neurons (Fig. 7g). Incorporating hold-duration predictive start-neuron assembly activity into behavioural models (Fig. 7h) improved prediction of trial-by-trial adjustments beyond reward history alone (Fig. 7i,j). This showed that start-neuron assembly activity accounted for variability between trials with identical reward outcomes but followed by different duration changes (for example, correct trials followed by an increase of 0.1 s versus 0.5 s). Thus, trial-onset activity within a cross-regional assembly of start neurons established the initial population state and predicted trial-by-trial behavioural adjustments.

## Discussion

In this study, we examined how internal estimates of elapsed time are represented across neural populations and how these representations relate to flexible timing behaviour. In a self-paced reaching task, mice generated accurately timed responses by adjusting hold durations based on trial history, in the absence of instructing sensory cues. Neural activity across all recorded regions decoded elapsed time in a relative reference frame. Despite heterogeneous single-neuron responses, distributed activity was organized within a low-dimensional ring manifold. Within this shared manifold, population activity evolved along a common trajectory across trials, with hold duration represented by traversal speed rather than distinct states (Fig. 8). These population dynamics were associated with coordinated co-scaling of single-neuron responses, with individual neurons jointly modulating, compressing or dilating their activity patterns in proportion to the interval. Activity of a cross-regional assembly of start neurons predicted the trial-by-trial variability in timing behaviour already at trial onset, linking initial population states to trajectory evolution.

**Fig. 8:**
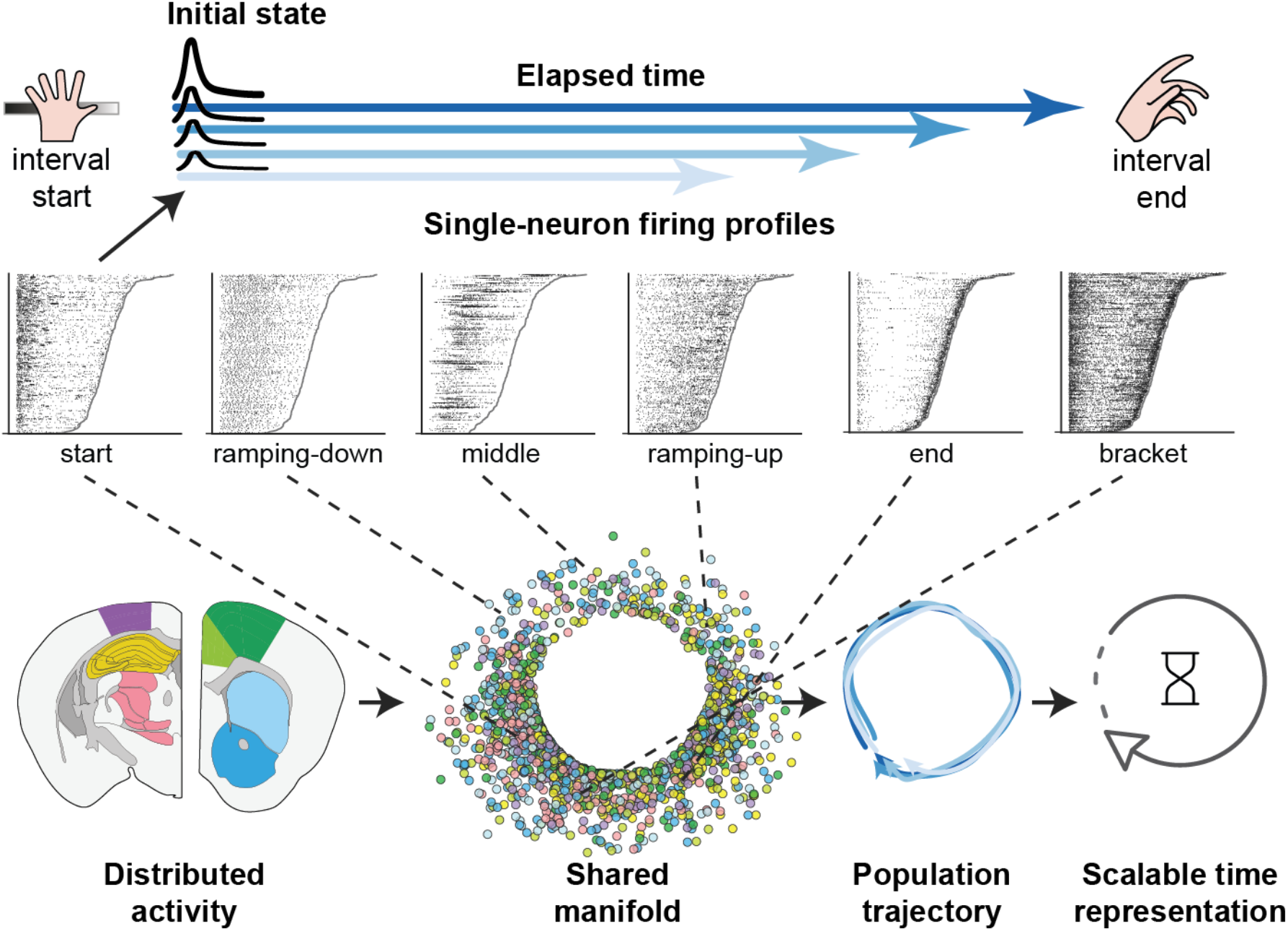
Summary schematic. In a self-paced interval timing task, accurate behaviour relied on an internal representation of elapsed time. Single neurons across brain regions exhibited heterogeneous temporal response profiles (e.g., start, ramping, middle, end, bracket), with trial-onset activity of a cross-regional assembly of start neurons setting the initial population state and predicting upcoming interval duration. At the population level, this heterogeneity was organized within a shared cross-regional manifold, in which angular position reflected each neuron’s preferred time within the interval. Population activity evolved along a common trajectory on this manifold across trials, with traversal speed encoding interval duration, supporting a scalable and adaptive representation of time.

In our self-paced interval timing task, mice generated reaching movements aligned to the target delay and adjusted hold durations based on recent outcomes. This is consistent with behaviour reflecting internal estimates of elapsed time rather than cue-driven responses or random trial-to-trial variability. Initiation and termination of the holding interval were self-determined, and the target delay was not instructed by cues but acquired through trial-and-error learning. Behavioural variability therefore more directly reflected fluctuations in internal time estimates, enabling to relate trial-by-trial variability in behaviour to population dynamics.

Task-related neural activity during the hold duration was distributed across all recorded regions. These responses exhibited structured heterogeneity, with diverse temporal firing profiles supporting decoding of elapsed time throughout the interval. Whether time is represented in relative or absolute coordinates in the brain remains uncertain^37^. In our self-paced task, temporal information was preferentially encoded in a relative rather than absolute reference frame. This relative representation supports generalization across durations by allowing rescaling of the shared population trajectories without requiring distinct neuronal ensembles or firing patterns^26^.

Single-neuron responses were highly consistent across individual trials, and included both monotonic and non-monotonic profiles, as previously reported^41^. Important distinctions, however, are worth noting. Specifically, middle neurons fired at specific times within the interval, and their response typically scaled with trial duration. In this respect, these middle neurons differ from early reports of time-cells, which are predominantly anchored to absolute time (e.g., firing at 2 s) and formed qualitatively new representations rather than scaling^8,42,43^. Middle neurons thus more closely resemble scaling time cells reported in later studies^34,40^. In fixed-interval task designs, however, the scaling might not have been directly observable, so neurons with similar properties may have been previously classified as time cells anchored to absolute time. Finally, middle neurons in our dataset did not fully tile the interval, even within individual regions, and also included sustained responses. This may reflect the nature of our task, where animals remained stationary during the hold duration, unlike most studies where time cells, and sequences in general, have been observed^8,42,44,45^ (but see also^46^) . Also notable, task-bracketing neurons have been previously described in the dorsolateral striatum and proposed to mark the boundaries of chunked action sequences ^47^, but with subsequent work pointing to a role beyond action execution ^48^. Here, bracket neurons were observed for the first time across multiple brain regions in the context of self-paced interval timing.

At the population level, the heterogeneity observed across single-neuron responses was captured by a ring-shaped manifold, with neurons arranged along a continuum and in which the angular dimension reflects each neuron’s preferred firing time within the trial (Fig. 4). This organization contrasts with population models in which time intervals are represented by sequential Gaussian-like time cells that tile the interval. In such cases, neurons with localized, time-locked responses would give rise to a linear manifold, where position along the line corresponds to preferred firing time (from start to end of the interval). In our data, the presence of bracket neurons, which fire at both the start and end of the interval, effectively links the boundaries of the sequence, allowing this linear structure to wrap into a circular manifold. Similar low-dimensional, non-linear manifolds have been observed in the entorhinal system, where grid-cell population activity forms a toroidal geometry, rather than a ring^25^.

This structure provides a geometric organization of diverse firing profiles across regions into a coherent representation of elapsed time. The results align with intrinsic models of timing predicting that elapsed time is encoded by evolving population dynamics rather than by dedicated clocks^2,49–51^. However, contributions to the common timing process were not uniform across regions despite being distributed: prefrontal and motor cortices showed higher decoding accuracy, whereas thalamus showed stronger coordination of decoding errors. Moreover, the prevalence of distinct single-neuron response profiles varied across regions, indicating non redundant contributions to the low-dimensional manifold ring. Interval timing therefore results from distributed dynamics with region-specific weighting rather than a homogeneous network or centralised mechanism. Population activity did not form distinct patterns across trials with different durations but progressed along a shared trajectory whose traversal speed scaled with hold duration. This contrasts with behaviour in which alternative choices are associated with distinct population states or divergent trajectories ^43,44,52^. This organization is consistent with population clock frameworks^49,53^, in which elapsed time corresponds to the evolving state of population activity along stable trajectories. Within this framework, time is encoded by position along this trajectory, and hold duration is determined by its traversal speed. The present findings provide network-level evidence for this principle across multiple brain regions and show that scalable traversal of a shared manifold accompanies flexible behavioural timing.

The coordinated cross-regional co-scaling uncovered during self-paced interval timing reveals that temporal scaling is a population-level property arising from structured trial-by-trial adjustments across neurons and regions, rather than from independent changes in isolated cells or trials. Co-scaling relationships were present across all recorded regions, pointing to a network-level phenomenon rather a local circuit effect. This distributed coordination provides a mechanistic account of how heterogeneous temporal response profiles can collectively generate stable yet flexible representations of elapsed time. By preserving the geometric structure of the shared manifold while modulating traversal speed, coordinated population co-scaling maintains coherent neuronal dynamics across trials with different durations. These findings extend prior accounts of scaling within individual regions^11,12,26,34,40,54,55^ to a brain-wide network.

A cross-regional assembly of coactive neurons exhibited early activity predictive of trial-by-trial timing behaviour. Similar predictive signals have been reported in motor cortex during self-initiated decisions^56^; here, these predictive dynamics were observed across brain regions in a paradigm with minimal movement confounds. This distributed organization indicates that predictive signals linked to internally guided behaviour reflect a general and network-level property rather than a feature of motor areas. Activity at trial onset within this assembly of coactive start neurons predicted behavioural timing variability beyond reward history and hence distinguished trials with identical outcomes but different hold duration history. These signals determined the initial state of population activity for each trial, and may reflect latent internal variables, such as expectation or recent experience, that influence subsequent population dynamics and trajectory speed. This suggests a role for this assembly in tracking task-relevant internal variables linked to interval timing on a trial-by-trial basis, rather than reflecting slower state changes.

Together, these results show that interval timing arises from heterogeneous, distributed, and yet geometrically organized population dynamics. During time estimation, population activity evolves along a stable low-dimensional trajectory on a shared manifold, with traversal speed determining hold duration. Temporal scaling is expressed at the level of single neurons but emerges as a coordinated, cross-regional process. An assembly of coactive start neurons predicts, at trial onset, the trajectory dynamics and timing behaviour of the ongoing trial. The distributed nature of these dynamics raises questions about the circuit mechanisms underlying cross-regional coordination. The presence of temporal information across multiple regions, coordinated decoding errors, and population co-scaling suggest structured interactions across regions, potentially mediated by recurrent connectivity, shared inputs, or modulatory signals^30,31,57–61^. In such a distributed system, perturbations at individual nodes could influence the rest of the network and reshape global dynamics, complicating the interpretation of region-specific manipulations.^27,62–66^ These findings establish cross-regional coordination as a central feature of brain representation for time and motivate future work to determine how such interactions are implemented at the circuit level.

## Methods

### Animals

These experiments used adult male C57BL/6 mice (2–4 months; Charles River Laboratories, UK). Mice were housed with their littermates until the surgical procedure. All mice were held in individually ventilated cages, with wooden chew sticks and nestlets, in a dedicated housing facility with a 12/12 h light/dark cycle (lights on at 07:00), 19–23◦C ambient temperature and 40–70% humidity. Food and water were available ad libitum unless stated otherwise. All experiments were conducted in accordance with the UK Animals (Scientific Procedures) Act 1986 under personal and project licenses issued by the UK Home Office following ethical review.

### Surgical procedure

Mice were implanted with a titanium headplate (Get It Made Ltd, London, UK) under isoflurane anaesthesia (4% induction, 1.5–2% maintenance). Analgesia was provided with buprenorphine (Vetergesic, 0.08 mg/kg, subcutaneous) and local anaesthetic (bupivacaine, 6 mg/kg) administered before skin incision. The scalp over the dorsal skull was removed and the surrounding skin secured to the skull using tissue adhesive (Vetbond, 3M, USA). Bregma was identified and stereotactic coordinates were used to mark target regions for subsequent craniotomies. The headplate was then secured on the skull surface using dental cement (Super Bond C&B, Parkell). A reference screw, wrapped in silver wire, was implanted over the cerebellum and fixed with dental cement (Jet Denture Repair Powder, Lang Dental, Illinois, USA; Meadway Repair Liquid, MR. Dental, Surrey, UK). A custom-designed 3D-printed protective shield was attached to the headplate to prevent the animal from reaching recording probes. After implantation, mice recovered for at least a week before further procedures.

One day before recordings, craniotomies were performed under isoflurane anaesthesia (4% induction, 1.5–2% maintenance) with buprenorphine analgesia (Vetergesic, 0.08 mg/kg, subcutaneous). Craniotomies were drilled using a fine dental drill at previously marked coordinates (anterior: AP +0.9 mm, ML 1.0 mm; posterior: AP −2.1 mm, ML 1.5 mm, relative to bregma). The dura was left intact and the exposed surface was kept moist with sterile saline throughout the procedure. Craniotomies were sealed with silicone elastomer (DuraGel, Cambridge Neurotech) and a biocompatible silicone (Body Double, Smooth-On) to protect the tissue and prevent dehydration until recordings.

### Behavioural task

This study involves a timed self-paced reaching task. Mice were habituated to head fixation and the experimental setup over up to five sessions, with fixation duration gradually increased from a few minutes to ∼25 minutes. Following habituation, mice were trained to perform left forelimb reaching movements toward a spout to obtain water droplets (4–5 μl, 5% sucrose)^67,68^. Water access was controlled during training to maintain body weight at ∼90% of pre-surgical weight (always >85%), with a minimum of 1 ml water provided daily.

Training proceeded in stages. Initially, mice were encouraged to perform left forelimb movements by applying a small droplet to the left whisker pad, paired with an auditory cue (3.6 kHz, 80 ms) and reward delivery at the spout. The spout was initially positioned close to the mouth and was gradually moved beyond tongue reach to promote forelimb reaching and discourage licking. Reaching movement toward the spout, detected with an infrared sensor (Panasonic Industrial Automation, FX-301HP), initially triggered reward delivery on every trial. After acquisition of reaching behaviour toward water droplets (position: ∼-0.5 mm anterior, ∼5 mm ventral, ∼5 mm lateral to the nose tip), mice were trained in the self-paced task requiring them to hold a metal handbar with the left forepaw before initiating a reach. Two self-generated cues were present during the task: a first tone (2.5 kHz, 80 ms) was triggered when mice started handbar holding that started the timer, and a different one (3,6 kHz, 80 ms) when their reaching movement was detected by the sensor. A target delay was set for each session. Premature bar release reset the timer. Reaches performed before the target delay triggered the second tone but no reward, whereas reaches performed after the target delay resulted in droplet delivery. Mice reached criterion performance within 1–2 weeks and performed >100 reaches per 30 min session. Following stable performance, mice underwent craniotomy and electrophysiological recordings (one session per day for 4–6 days). Only sessions with target delays >1 s were included in analyses (56 sessions from 9 mice).

### Behavioural setup

Behavioural training and recordings were performed in a custom-built head-fixed rig constructed from Thorlabs components and inspired by the International Brain Laboratory design^69^. A 3D-printed headplate holder (Rigid 10k resin, Formlabs) was used to secure the animal. Task control and event timing were implemented using the pyControl system^70^. Bar contact and spout contact were detected electrically using a lickometer circuit combined with a dual comparator lick port detector (Janelia Experimental Technologies) to minimize electrical artifacts during recordings. Reaching movements were detected using an infrared sensor (FX-301HP, Panasonic Industrial Automation) and used to compute holding time and trigger reward delivery in case of correct trials. Water droplets (4–5 µl) were delivered via a gravity-fed system controlled by a miniature solenoid valve (LFVA1220210H, Lee Company). Auditory cues were delivered through a speaker positioned ∼10 cm from the animal. Behavioural events (for example, handbar touch, handbar release, spout contact) were timestamped using pyControl and used to trigger task events in real time. Video recordings were acquired from front and side views at 100 frames per second using high-speed near-infrared cameras (MQ013RG-ON, Ximea or PL-D721P, Pixelink). Two LED illuminators (BW 48 LED) were used for infrared lighting. Camera frames were triggered at 100 Hz by the pyControl system to ensure synchronization with behavioural events. Video data were acquired using StreamPix9 (Norpix) and encoded in H.264 format.

### Electrophysiology

Recordings were obtained from 6 of 9 mice (31 sessions) trained in the timed self-paced reaching task using Neuropixels 1.0 probes. Two probes were used per session (referred to as probe 1 and probe 2). Probe 1 targeted prefrontal cortex (anterior cingulate, infralimbic and prelimbic cortex; ACA, ILA, PL), motor cortex (primary and secondary motor cortex; MOp, MOs), dorsal striatum (caudate putamen; CP) and nucleus accumbens (NAc). Probe 2 targeted parietal cortex (PPC), hippocampus (Hpc) and posterior thalamus (PO, LP, LD, VM, VPM, VPL). The choice of these regions was guided by prior studies examining timing-related activity in individual regions in isolation^4,7–9,29,71,72^

Probes were mechanically sharpened and the ground and reference pads were soldered to a common ground wire connected to the animal. Before each insertion, probes were coated with fluorescent dye (DiI, 1mg/ml in isopropanol) for post hoc track reconstruction.

Probes were mounted on linear motor drives (IVM, Scientifica) attached to stereotaxic manipulators (Kopf Instruments 1363A, 3108B). Insertions were performed simultaneously at 3 µm/s (probe 1: depth 4.8 mm, 3–5° mediolateral tilt; probe 2: depth ∼2.9 mm). After reaching target depth, probes were retracted by 100 µm and allowed to settle for 5–10 min before recording. Recording sessions consisted of a behavioural training block (30 min) preceded and followed by baseline epochs (up to 20 min each). After each session, probes were retracted at a speed of 5 µm/s and cleaned using enzymatic detergent (Terg-a-zyme, 1%, Sigma Aldrich), followed by isopropanol and distilled water.

Signals were acquired using OpenEphys in binary format. Default acquisition settings were used (AP band: 30 kHz sampling rate, 500× gain; LFP band: 2.5 kHz sampling rate, 250× gain; 384 channels per probe). Recordings were referenced to the probe tip. Electrophysiological data were synchronized with behavioural events and video frames using TTL pulses generated by pyControl. The Rsync module of pyControl was used to synchronize the electrophysiological data, behavioural events and the triggered video frames for downstream analyses.

### Histology and imaging

At the end of all recording experiments, mice were anesthetized with a terminal intraperitoneal injection of pentobarbital (200 mg/kg) and transcardially perfused with 0.1 M phosphate buffered saline (PBS), followed by 4 % paraformaldehyde solution (PFA) in PB. Brains were extracted and post-fixed overnight in 4 % PFA in PB at 4°C in the dark, then transferred to 30 % sucrose in 0.1 M PB for cryoprotection at 4°C overnight. Brains were sectioned coronally at 50 µm using a freezing microtome (Epredia HM 450) and stored in PB containing sodium azide until mounting for imaging. Fluorescent DiI tracks and brightfield images were acquired using the Cy3 imaging protocol (Zen Blue software, excitation using 532 nm laser line) on an epifluorescence microscope (Axio Imager M2, Zeiss) with a 5× objective.

### Region assignment

Probe tracks were reconstructed using the SHARP-Track toolbox^73^. Histological sections were aligned to the Allen Mouse Brain Common Coordinate Framework (CCF), and probe trajectories were manually traced based on DiI fluorescence. Individual probe insertions were assigned to recording sessions by matching reconstructed trajectories to experimentally documented entry coordinates (manipulator positions). When fewer tracks were recovered than insertions, the closest matching trajectory was assigned. Channel locations along each probe were mapped to CCF region labels using adapted SHARP-Track code.

Region boundaries were further refined using electrophysiological landmarks along the probe. A graphical user interface^68^ was used to visualize spiking and local field potential (LFP) features across channels, including: spike raster, firing rate, cell count, waveform peak amplitude, multi-unit spiking activity (number of crossing below −100 µV in 30 s interval), raw LFP (after subtracting median across channels), LFP power (Welch’s power spectral density estimate), LFP power normalized across channels, LFP power relative change across channels, LFP cross correlation across channels. We visually inspected each probe insertion and shifted the probe to align region boundaries to characteristic electrophysiological landmarks^68,74^.

### Spike sorting and quality control

Automated spike sorting was performed using KiloSort 3.0. with default parameters^75,76^. A total of 52,242 clusters were initially identified, of which 24,051 (46%) were classified as “good” by Kilosort and retained for further analysis. Units were imported into the CellExplorer framework and Phy for visual inspection of event-aligned spike raster, waveform, and auto-correlograms. Additional curation was performed using the Bombcell toolbox^77^ with parameters detailed Table S2, yielding 11,050 units (21.1%). For analyses involving correlations or distances between population activity vectors, units were further assessed based on mean firing rate, retaining only neurons with an average firing rate of at least 0.1 Hz across the entire recording session (final dataset: 8,642 units). For analyses involving trial-wise regressions, trials were restricted to durations between 1 and 10 s to mitigate the impact from short or long trials, unless stated otherwise.

### Distribution and cross-session evolution of hold intervals

To characterize holding behaviour, we analysed the distribution of hold intervals (seconds) across all trials for each session and mouse (e.g., Fig. 1d). To account for differences in session-specific target delays, hold intervals were normalised as:

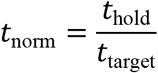

such that *t*_norm_ = 1 corresponds to the session target delay (e.g., *t*_norm_ = 1.2 indicates a hold 20% longer than the target delay; see Fig. 1d).

For each session and mouse, we computed the empirical cumulative distribution function (CDF) of *t*_norm_ to quantify behavioural pacing within and across sessions. Session-wise CDFs were fitted with a three-parameter logistic function using the curve-optimization package from SciPy:

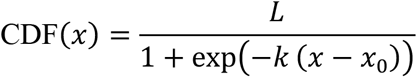

where *x* is normalized hold time, and *L*, *k*, and *x*_0_ correspond to the asymptote, slope, and inflection point of the fit, respectively. Fitted parameters were retained for each session and mouse for subsequent analyses.

To assess changes in performance across sessions (Fig. S1d), we quantified the relationship between session index and the slope parameter *k*. For each mouse, Spearman’s rank correlation was computed between session-wise index and *k* values across sessions. To estimate chance levels, session indices were circularly permuted 1,000 times, and the Spearman correlation recomputed for each permutation to generate a null distribution against which observed values were compared.

### Trial-history regression analysis

To determine whether behaviour was influenced by recent trial outcomes, we implemented linear regression models (Fig. 1e,f). For each session, categorical predictor variables encoded the outcomes of the three preceding trials (trials *t* − 1, t − 2, and t − 3). Trial outcomes were represented as binary variables (premature = 1, rewarded = 0). The dependent variable was defined as the change in hold duration between consecutive trials:

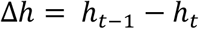

where ℎ*_t_*_–1_ is the hold time in trial t −1, and ℎ*_t_* is the hold time in trial t.

Models were fitted separately for each session containing at least 100 trials. Data were split into training (80%) and test (20%) sets. Model performance was quantified as the Pearson correlation between observed and predicted changes in hold duration in the test set. This cross-validation procedure was repeated 20 times with different random splits of the data, and model accuracy was defined as the mean correlation across repetitions.

To control for regression-to-the-mean effects (i.e., prediction driven by the distribution of hold durations rather than trial outcome), we generated a null distribution by permuting the dependent variable Δℎ values 500 times and refitting the model on each permutation. The resulting surrogate accuracy distribution was used to z-score the observed model accuracy, yielding a normalized prediction accuracy. A schematic of this analysis is shown in Fig. 1e, and the corresponding results are reported in Fig. 1f and Fig.S1e.

### Off-task control epochs

Pre- and post-task periods included off-task control epochs during which mice were not engaged in interval timing task but could hold the handbar (Fig. S2a). These epochs were used as control periods to assess whether observed effects were specific to task engagement rather than reflecting passive or temporally unstructured behaviour. For each session, off-task control intervals were sampled from these periods to match the number and duration distribution of hold intervals observed during the task. This procedure was applied in the analyses shown in Figs. 2–4 and 7.

### Peri-event time histograms

Peri-event time histograms (PETHs) were constructed by aligning spike times to either handbar touch or handbar release (Fig. 2b). PETHs were computed within an 8-s window centred on the trigger event (±4 s) using 200-ms bins. To avoid contamination from adjacent trials, analyses in Fig. 2b,c included only trials for which the preceding reach occurred at least 2.5 s before the next trial onset. As an additional control for potential carry-over effects, PETHs were recomputed using only trials not preceded by a rewarded trial (Fig. S2c). Control PETHs were also computed during off-task windows (see section “Off-task control epochs”) using the same alignment and binning procedures. For analyses in Fig. S4f, Fig. 7a and Fig. 6a, PETHs were computed separately for different ranks of trial duration to compare neuronal responses across trials of varying lengths.

### Extracting dominant population responses using PCA

Trial-averaged peri-event time histograms (PETHs) were computed for each neuron by aligning spike times to handbar touch or release (Fig. 2b). To characterize the heterogeneous response profiles observed across individual neurons, for each session principal component analysis (PCA) was applied to the population of PETHs to capture dominant patterns of activity, and the first three principal components (PCs) were retained. After verifying that the dominant PCs were consistent across sessions and mice, we computed a common PC space using the full dataset and projected all neuronal responses onto this shared subspace.

The projections within this PC space were used to quantify neuronal modulation strength. For each neuron, modulation strength was defined as the Euclidean norm (vector length) of its coordinates in the PC space. Session-level modulation was computed as the mean across neurons (Fig. 2d). As a control, the same procedure was applied to PETHs derived from off-task epochs.

To assess whether neuronal responses across brain regions occupied overlapping or distinct subspaces within this PC space, we trained a linear discriminant analysis (LDA) classifier to predict region identity from the PC projections. To control for class imbalance, neurons were subsampled such that each region contributed an equal number of units, determined by the minimum number recorded in any region. Classification performance was quantified using recall scores for each region (Fig. 2f). Chance-level performance was estimated by repeating the analysis in surrogate LDA models with randomly shuffled region labels, thereby disrupting the relationship between neuronal response profiles and anatomical identity and generating a null distribution of recall scores. This procedure was repeated 1,000 times with independent subsampling of neurons from each region. As an additional control, the same analysis was performed using PETHs during off-task windows (Fig. S2e).

### Single-neuron response heterogeneity

To assess the similarity of neuronal responses within and across anatomical regions, we quantified response heterogeneity based on PETHs (see section “Peri-event time histograms”). For each session, we computed the pairwise cosine distance between single-neuron response profiles, using the PETH-derived response vectors aligned to handbar touch and handbar release. This yielded an *N* × *N* distance matrix for each session, where N denotes the total number of recorded neurons. We quantified heterogeneity by comparing the mean cosine distance between neuron pairs within the same region (within-region distance) to that between neuron pairs from different regions (across-region distance). Higher cosine distance indicates greater dissimilarity in response profiles.

### Elapsed time regression models

To assess whether neural population activity encoded elapsed time between handbar touch and handbar release, spike counts were computed in 200-ms bins for trials lasting at least 1 s (Fig. 3). Neural activity within the 200 ms preceding handbar release was excluded to minimize contamination from movement-related signals.

Elapsed time was decoded from population activity using a k-nearest neighbours (kNN) regressor (k = 5; Euclidean distance; scikit-learn), allowing for non-monotonic (e.g., Gaussian-like) temporal tuning profiles. For each session, trials were randomly split into training (80%) and testing (20%) sets. Neural activity was z-scored using the mean and standard deviation computed from the training set and applied to both training and testing data. Model performance was evaluated as the Pearson correlation between actual and predicted elapsed time in the test set. This cross-validation procedure was repeated 20 times with different random splits, and session-level accuracy was defined as the mean correlation across repetitions. Chance-level performance was estimated by circularly shuffling elapsed time labels and repeating the full training and testing procedure. Observed accuracy was z-scored relative to this null distribution to obtain a chance-normalized decoding accuracy.

Two encoding schemes were tested. In the absolute time model, elapsed time corresponded to the physical time since handbar touch (e.g., 1 s from trial onset). In the relative time model, elapsed time was expressed as the fraction of total trial duration (e.g., 10% of the interval), thereby normalizing across intervals of different lengths. As an additional control, identical analyses were performed on duration-matched off-task intervals (see section “Off-task control epochs”) to assess whether elapsed time encoding was specific to task engagement (Fig. 3d,e).

To quantify the relative contribution of absolute and relative time coding at the single-neuron level (Fig. S3a,b), regression analyses were performed for each neuron individually. For each neuron, spike counts were computed in 200 ms bins, and regression models were trained separately to predict binned activity from either absolute or relative elapsed time. Model performance was quantified as the Pearson correlation between observed and predicted spike counts. To estimate chance levels, time labels (absolute or relative) were shuffled before model fitting, generating surrogate accuracy distributions. Observed performance was then z-scored relative to these distributions to obtain a chance-corrected measure of encoding strength. Each neuron was thus characterized by two values: absolute time encoding (with relative time labels shuffled) and relative time encoding (with absolute time labels shuffled).

### Regional and ring-angle elapsed time models

To assess how relative elapsed time encoding varied across anatomical regions (Fig. 3f), regression models were trained separately for each region. To control for differences in neuron counts across regions, a fixed number of neurons (n = 10) was randomly sampled from each region to form the population input to the model. This subsampling procedure was repeated 250 times per region. Regions with fewer than 10 neurons in a given session were excluded from that session’s analysis.

An analogous analysis was performed using groups of neurons defined by angular sectors of the ring manifold rather than anatomical region (Fig. S4e; see Section ‘Ring manifold’). For each sector, 10 neurons were randomly sampled and used to train regression models as described above. This approach enabled comparison of elapsed time encoding across neurons grouped by firing profile class rather than anatomical location.

### Decoding error correlations across regions

To assess whether trial-by-trial fluctuations in elapsed time decoding were coordinated across regions (Figs. 3g and S3c,d), we quantified correlations in decoding errors between regions. For each session, we obtained trial-level elapsed time predictions using region-specific regression models (see “Regional and ring-angle elapsed time models”). For each region and trial, decoding error was defined as the mean squared error (MSE) between predicted and actual elapsed time, yielding, for each session, a vector of trial-wise decoding errors per region. Pairwise Pearson correlations were computed between the trial-level decoding error vectors of all region pairs, resulting in a symmetric region-by-region correlation matrix (7 × 7) representing the extent to which decoding errors co-fluctuated across regions. The mean of the off-diagonal elements was taken as the average cross-regional decoding error correlation for that session.

To estimate chance levels, we generated 500 surrogate datasets in which decoding error vectors were independently circularly shuffled within each region, thereby disrupting trial-wise alignment across regions while preserving each region’s error distribution and temporal structure. The full procedure was repeated for each surrogate dataset to obtain a null distribution of mean cross-regional decoding error correlations.

### Extracting warped-time tuning curves

Warped-time tuning curves were computed for individual neurons by expressing spike counts as a function of relative elapsed time between handbar touch and release (see “Elapsed time regression models”; Fig. 4a). This allowed aligning neural activity across trials of different lengths through a time-warping transformation. To construct tuning curves, relative elapsed time was discretized into 10 equally spaced bins (unless stated otherwise). For each bin, we computed the mean firing rate across trials, yielding a tuning curve that describes the average response profile of the neuron as a function of warped time. Tuning curves were z-scored independently for each neuron prior to further analyses.

### Ring manifold

To characterize the diversity of neuronal response profiles (see “Extracting warped-time tuning curves”), we applied non-linear dimensionality reduction to warped-time tuning curves. Specifically, Isomap (scikit-learn manifold module; 30 neighbours; cosine distance metric) was used to project the 10-dimensional tuning curves of all neurons into a low-dimensional manifold space.

The resulting manifold exhibited a ring-like geometry (Fig. 4b). To characterise this structure, we fitted a circle to the two-dimensional projection using least-squares optimization (scipy.optimize), according to:

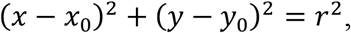

where (*x*_0_, *y*_0_) denotes the centre and *r* the radius of the ring. The manifold was then re-centred using the fitted centre, and each neuron was represented in polar coordinates defined by its angular position and radial distance. As a control, the same procedure was applied to tuning curves derived from off-task epochs (see “Off-task control epochs”).

To assess the stability of ring positions across trials, we performed a cross-validation analysis. For each session, tuning curves were computed using 50% of trials (training set), and the ring was fit from this subset as described above. Tuning curves from the remaining 50% of trials (test set) were projected onto the training-derived manifold, and each neuron’s angular position within the ring was estimated. For each neuron, we computed the angular difference between training- and test-derived positions. Small angular differences indicate stable response profiles across trials, reflected by consistent ring positions. This procedure was repeated 1,000 times with random splits of trials into training and test sets, and the mean angular difference across permutations was used as a measure of stability. As a control, the same cross-validation procedure was applied to off-task tuning curves. A schematic of this approach is shown in Fig. S4c, and the corresponding results are reported in Fig. S4d.

For subsequent analyses, a final manifold was constructed using tuning curves from all trials. Based on subsequent analyses supporting the ring topology (see section ‘Persistent homology analysis of ring manifold), only the first 2 dimensions of the manifold were retained.

### Supervised extraction of single-neuron response profiles

Single-neuron response profiles were classified into six categories: start, ramp-up, ramp-down, middle, end, and bracket neurons (Fig. 4a). To relate these profiles to the ring manifold, template tuning curves corresponding to each response type were constructed (Fig. 4d). Templates were defined over 10 bins of relative elapsed time, consistent with the warped-time tuning curves. Start templates were defined by high activity (value = 1) confined to the first bin and zero elsewhere. End templates by high activity in the final bin and zero elsewhere. Bracket templates by high activity in both the first and final bins. Middle templates by high activity in the central three bins and zero elsewhere. Ramp-up and ramp-down templates were defined as monotonically increasing and decreasing profiles, respectively, across bins. For each neuron, the Pearson correlation between its warped-time tuning curve and each template was computed, and the neuron was assigned to the class yielding the highest correlation.

For the analyses presented from Fig. 4d onwards, we mapped template-defined response classes onto the ring manifold. After assigning each neuron a template label, we trained a linear discriminant analysis (LDA) classifier to predict template class from the ring-centred angular coordinate. The trained model was then used to define the angular sectors corresponding to each response profile (see Fig. 4d captions). This approach was used to partition neurons in Fig. 4d and to order neurons by angular position (relative to start neurons) in Fig. 5a.

### Scaling index

To quantify the consistency of neuronal responses across trials of different durations, we computed a scaling index metric adapted previous work^26^. For each session, trials were sorted by hold duration and divided into quartiles. For each neuron, warped-time tuning curves (see “Extracting warped-time tuning curves”) were computed separately for each duration quartile. Pairwise Pearson correlations were then calculated between quartile-specific tuning curves, and the mean correlation was defined as the scaling index for that neuron. Higher values indicate greater similarity of the response profile across different interval durations, consistent with temporal scaling.

To estimate chance levels, we generated 1,000 surrogate datasets by independently shuffling time bins within each quartile-specific tuning curve, thereby disrupting structured temporal alignment. A scaling index was computed for each surrogate, yielding a null distribution. P values were obtained by comparing observed values to this distribution for each neuron. Neurons with *p* < 0.05 were classified as significantly scaling, indicating consistent response profiles across trials of different durations.

### Persistent homology analysis of ring manifold

To formally characterize the geometry of the manifold (e.g., whether it resembled a cluster, a ring, or a higher-dimensional structure), we applied persistent homology using the Ripser package (ripser.py; https://github.com/scikit-tda/ripser.py/). Persistent homology quantifies the presence and stability of topological features in a point cloud as a function of scale. A radius is progressively increased around each point (i.e., a neuron in the ring), and topological features that persist across scales are identified. These features are summarized by homology groups, commonly denoted *H*_0_, *H*_1_, and *H*_2_. *H*_0_ corresponds to connected components, *H*_1_1 to loops, and *H*_2_ to enclosed cavities. The persistence of these features can be visualized as barcodes (Fig. S4a-b), where longer bars indicate features that persist across a broader range of scales.

Analyses were performed on manifolds derived from task-related tuning curves and from off-task control data (see “Ring manifold”). To reduce computational cost, neurons were subsampled across the manifold by selecting 20 neurons within each 45° angular bin. To limit the influence of outliers in the manifold, this analysis was restricted to neurons exhibiting significant tuning-curve consistency across trials of different durations (see “Scaling index”). Persistent homology was computed for each subsample. Topological features were identified from barcode representations of homology groups (*H*_0_, *H*_1_, *H*_2_). Within each subsample, a feature was considered present if at least one barcode length exceeded 10× the mean barcode length for that subsample. This procedure was repeated 1,000 times to estimate the occurrence of connected components (H₀), loops (H₁), and cavities (H₂) in the manifold.

### Structure index of ring manifold

To determine which latent variables best accounted for the geometry of the ring manifold, we computed the Structure Index^39^. This metric quantifies the extent to which distances in an N-dimensional manifold (here, the manifold defined by neuronal tuning profiles) can be explained by an external variable. For each neuron, we considered the following candidate variables: (1) preferred firing time within the interval, defined as the peak bin of the warped-time tuning curve; (2) scaling index (see section “Scaling index”); (3) anatomical region; (4) mean firing rate during the task; (5) mean firing rate across the full session; and (6) putative cell type (principal cell versus interneuron).

For each session, we computed the Structure Index of each variable with respect to the ring manifold. Chance levels were estimated by shuffling variable values across neurons (250 repetitions), preserving their distribution while disrupting the relationship with manifold position. Observed values were then normalized relative to this null distribution, yielding a chance-normalized measure of how strongly each latent variable structured the ring geometry.

### Elapsed-time prediction from low-dimensional angular trajectory

To assess whether angular position in the ring manifold encoded relative elapsed time, each neuron’s angular coordinate was represented in two-dimensional Euclidean space as (cosθ, sinθ). For each session, instantaneous population spike-count vectors were computed in time bins as described in “Elapsed time regression models.” For each time bin, the instantaneous population angle was calculated as the activity-weighted mean of the neurons’ angular coordinates, using their spike counts in that bin as weights. Repeating this procedure across time bins yielded a time-resolved angular trajectory for each trial. A schematic and validation of this method is shown in Fig. S5a.

To determine whether these trajectories encoded elapsed time, and because the relationship between angle and elapsed time within a trial was non-linear (see Figs. 4 and 5a), we used a support vector machine (SVM) classifier with a nonlinear kernel to predict discretized relative elapsed time from the two-dimensional angular projection. Relative elapsed time was discretized into 10 bins (e.g., 0-0.1, 0.1-0.2, etc.). Models were trained using an 80:20 train-test split, and classification accuracy was quantified on held-out trials. This procedure was repeated 20 times with random train-test partitions, and mean accuracy was defined as the session-level decoding performance. Chance-level performance was estimated by generating 500 surrogate datasets in which the relative elapsed time labels were circularly shifted within each trial, thereby disrupting the alignment between neural activity and elapsed time while preserving the temporal structure of the activity and the distribution of time bins. The same training and testing procedure was applied to each surrogate dataset, yielding a null distribution of classification accuracies against which the observed decoding performance was compared.

To extract population trajectories (e.g. Fig. 5e), for each session, we grouped trials into 10 ranks based on trial duration. To enable comparison across trials of different durations, trajectories were first linearly resampled to 50 time points. Trials were then sorted by duration, assigned to ranks, and averaged within each rank. We quantified the traversal speed of each rank-averaged trajectory by computing the cumulative angular distance within each rank and dividing it by the mean trial duration of that rank (Fig. 5f). For each session, speeds were normalised by the value in the first rank (shortest trials). To assess whether normalised speed decreased with increasing rank, we computed the Spearman correlation between rank number and normalised speed and tested against zero. We quantified the similarity between rank-averaged trajectories by computing cosine similarity between trajectories and averaging these values within each session for three conditions: across duration ranks (similarity between different ranks, estimated using matched split-halves), within duration ranks (split-half similarity within each rank), and after temporal shuffling of rank-averaged trajectories (random permutation of temporal samples) (Fig. S5d).

To minimize the impact of sparse or noisy responses across trials, these analyses was restricted to neurons with a significant scaling index, indicating consistent firing patterns across intervals (see “Scaling index”).

### Population co-scaling analysis

To determine whether neurons scaled their activity in a coordinated manner across trials, rather than independently, we quantified trial-by-trial population co-scaling of neural responses. For each significantly scaling neuron (see “Scaling index”), spike counts were computed in 200-ms bins during the first second of each trial (aligned to handbar touch) and smoothed with a Gaussian kernel (s.d. = 200 ms). We restricted the analysis to the first second to minimize confounds from the overall trial duration. Only trials between 1 and 8 s in length were included to exclude excessively long trials associated with reduced task engagement.

To capture the dominant temporal response dynamics without imposing specific assumptions about scaling features (e.g., slope for ramping neurons or peak timing for middle/end/bracket neurons; see Fig. 6a), we applied principal component analysis to the z-scored waveforms of each neuron. The first principal component (PC1) was extracted, and each trial was assigned a projection score onto PC1. This score quantified how the primary temporal response pattern of each neuron varied across trials.

To measure co-scaling, we computed the trial-to-trial change in PC1 projection scores (i.e., difference between consecutive trials) for each neuron. For a given neuron, we then trained ridge regression models to predict its PC1 change from the PC1 changes of all other simultaneously recorded neurons. To control for collinearity, predictor neurons whose activity was highly correlated (*r* ≥ 0.85) with that of the target neuron were excluded. This approach tested whether changes in one neuron’s temporal dynamics could be predicted from concurrent changes in the population, indicating coordinated scaling. Models were trained using a 90:10 train-test split and repeated 10 times with different random splits. Prediction accuracy was quantified as the Pearson correlation between the observed and predicted PC1 changes in the held-out data. The mean accuracy across repetitions was taken as the neuron’s co-scaling accuracy.

To estimate chance levels, we generated 1,000 surrogate models in which predictor variables (i.e., PC1 change time series of other neurons) were circularly shuffled independently. This procedure preserved each neuron’s intrinsic scaling across trials while disrupting coordinated trial-by-trial fluctuations between neurons. The resulting surrogate accuracy distribution was used to compute numerical p values for each neuron, identifying neurons whose scaling dynamics were significantly coordinated with the rest of the neural population, i.e. significantly co-scaled.

In Fig. 6c, we report the session-level mean regression accuracy compared to the corresponding surrogate accuracy, as well as the number of significantly co-scaling neurons relative to the proportion expected under the null distribution. Finally, Fig. 6d shows region-wise regression coefficients derived from models of significant neurons, computed with respect to the full population.

### Ensemble dynamics and trial-duration prediction

To test whether ensemble activity associated with distinct response profiles (see section “Supervised extraction of response profiles”) predicted trial duration, we extracted handbar-touch–aligned spike activity from −3 to +6 s relative to touch and pre-processed as outlined in section ‘Population co-scaling analysis’. For each session, neurons were grouped according to their ring-defined response profile (start, ramp-up, ramp-down, middle, end, and bracket). Within each profile class, activity was averaged across neurons to obtain a session-specific ensemble response for each profile type (Fig. 7a). To assess whether ensemble activity predicted overall trial duration, we trained linear regression models using the mean activity of all ensemble profiles within a 0.6-s window centred on handbar touch as predictors. Predictor activity was z-scored prior to model fitting. Models were trained using an 80:20 train–test split and evaluated as described in the section “Population co-scaling analysis,” with accuracy quantified as the Pearson correlation between true and predicted trial duration in the held-out data. Chance-level performance and numerical p values were estimated by permuting trial-duration labels 1,000 times and repeating the full regression procedure to generate a null distribution of prediction accuracy.

To examine the temporal evolution of predictive information, we repeated this analysis using a sliding window (step size 200 ms), thereby estimating trial-duration prediction accuracy as a function of time relative to handbar touch. The resulting time-resolved decoding accuracy is shown in Fig. 7b, and the corresponding ensemble-specific regression coefficients are shown in Fig. 7c and Fig. S6b. This analysis used the same trial and neuron inclusion criteria as those highlighted in the section ‘Population co-scaling analysis’, restricting sessions to at least 100 trials and including only neurons with a significant scaling index (see section “Scaling index”).

### Hold-duration predictive start neurons

To identify start neurons whose activity at trial onset predicted whole-trial duration, we extracted spike counts as described in the section “Population co-scaling analysis.” For each trial, instantaneous firing rates were computed within the first 200 ms following handbar touch. For each start neuron, we then calculated the Spearman correlation between its early firing rate at trial onset and the corresponding whole-trial duration. The observed correlation was compared against a null distribution generated by 1,000 permutations of trial-duration labels. This procedure yielded, for each neuron, a correlation coefficient and associated p value reflecting its trial duration prediction. Start neurons with p < 0.05 were classified as significantly hold-duration predictive.

As a control group within each session, we selected a matched number of non-significant start neurons by ranking neurons in descending order of p value and selecting the same number (or as many as possible with p > 0.1) as the significant hold-duration predictive group. All analyses involving predictive start neurons were restricted to sessions meeting the same trial inclusion criteria described in the section “Population co-scaling analysis”. To avoid sparsely sampled sessions, analyses were further limited to sessions with at least two significantly predictive start neurons.

To further test whether start neurons formed hold-duration predictive assemblies, we quantified their millisecond-timescale coactivity during off-task control windows (see section “Off-task control epochs”). Off-task activity was used to avoid confounds related to within-trial covariation. Population spike-count vectors were computed in 50-ms bins. Pairwise Pearson correlations were then calculated between significantly predictive start neurons and, separately, between non-predictive start neurons.

### Behavioural regression model including neural activity

To assess whether trial-by-trial changes in timing behaviour (i.e., changes in hold duration) could be explained by neural activity in addition to recent trial outcome (Fig. 7h), we extended the behavioural regression model described in “Trial-history regression analysis” (Fig. 1e). Analyses focused on the most recent trial (*t* − 1), which exerted the strongest behavioural influence (Fig. S1e). The dependent variable was the change in trial duration between consecutive trials. Predictor variables included the outcome of the previous trial and the change in firing rate of start neurons between consecutive trials. Analyses were performed separately for hold-duration-predictive and non-predictive start neurons (see “Hold-duration predictive start neurons”). To quantify the contribution of trial-by-trial changes in start-neuron activity, we compared the model’s accuracy to that obtained from surrogate models in which trial outcome history was preserved but neuronal activity was shuffled. This procedure isolated the additional predictive contribution of neural activity beyond trial outcome in shaping behavioural adaptation on the subsequent trial. Models were trained separately for each session following the procedure described in the section “Behavioural trial-outcome model.” Sessions and trials included in this analysis followed the same criteria described in the section “Hold-duration predictive start neurons.”

### Statistical analysis

Data preprocessing was performed in MATLAB, and analyses were conducted in Python3.10 using NumPy^78^, pandas^79^, matplotlib^80^, seaborn^81^, SciPy^82^, scikit-learn^83^, NetworkX^84^, statsmodels^85^, and DABEST^86^. Two-sided statistical tests assuming symmetric distributions were visualized using Gardner–Altman and Cumming plots (DABEST), which report effect sizes as mean or median differences between groups (e.g., Fig. S1e). The upper panel shows raw data (mean ± SEM), and the lower panel shows bootstrapped differences (5,000 resamples) with mean and 95% confidence intervals. Comparisons against a fixed value or between two conditions were performed using bootstrap tests (paired or unpaired), estimating mean differences from 100,000 resamples with replacement. P values were computed numerically under the null hypothesis of no difference. For one-sided tests, p values correspond to the proportion of bootstrap samples exceeding zero in the specified direction; for two-sided tests, p values were obtained by doubling the smaller proportion of samples above or below zero. Unless otherwise stated, all p values are two-sided. Bootstrapped mean differences were also visualized as histograms (e.g. Fig. 7d,f,h), representing sampling distributions rather than raw data. Comparisons across multiple conditions were performed using linear mixed-effects models. For the mixed-effects analyses, condition (for example, brain region) was included as a fixed effect, and session was included as a random intercept to account for repeated measurements within sessions. Models were fit by maximum likelihood in statsmodels. To test the overall effect of condition, the full model (including condition) was compared with an intercept-only null model using a likelihood ratio test, and significance was assessed with the corresponding chi-square statistic. All confidence intervals (95% CI) were computed by bootstrapping (100,000 resamples, unless stated otherwise), using the 2.5th and 97.5th percentiles of the resampled distributions.

## Acknowledgements

We would like to thank B. Bathellier and W. Nicola for commenting on a previous version of the manuscript; B. Micklem, B. Perry, N. Shackle, and J. Westcott for technical assistance; all members of the Dupret lab for continued feedback during the project. This work was supported by the Medical Research Council (MRC) UK (MC_UU_12024/3, MC_UU_00003/4, and MR/W004860/1 to D.D. and MC_UU_00003/6 to A.S.). M.S. is supported by an SNSF Swiss Postdoctoral Fellowships (P500PB_206885/1). M.C. is supported by an MRC UK studentship (MR/W006731/1). Y.P. is supported by a German Research Foundation Walter Benjamin Fellowship (451242556).

## Author contributions

Conceptualization, M.S. and D.D.; Methodology, M.S. and D.D.; Data Collection, M.S.; Investigation, M.S.; Formal Analysis, M.S. and M.C.; Resources, M.S., M.C., Y.P., A.S., and D.D.; Writing – Original Draft, M.S., M.C., and D.D.; Writing – Reviewing & Editing, M.S., M.C., Y.P., A.S., and D.D.; Visualization, M.S., M.C., and D.D.; Supervision, D.D; Funding Acquisition, A.S., D.D.

## Declaration of interests

The authors declare no competing interests.

## Supplementary Figures

**Fig. S1:**
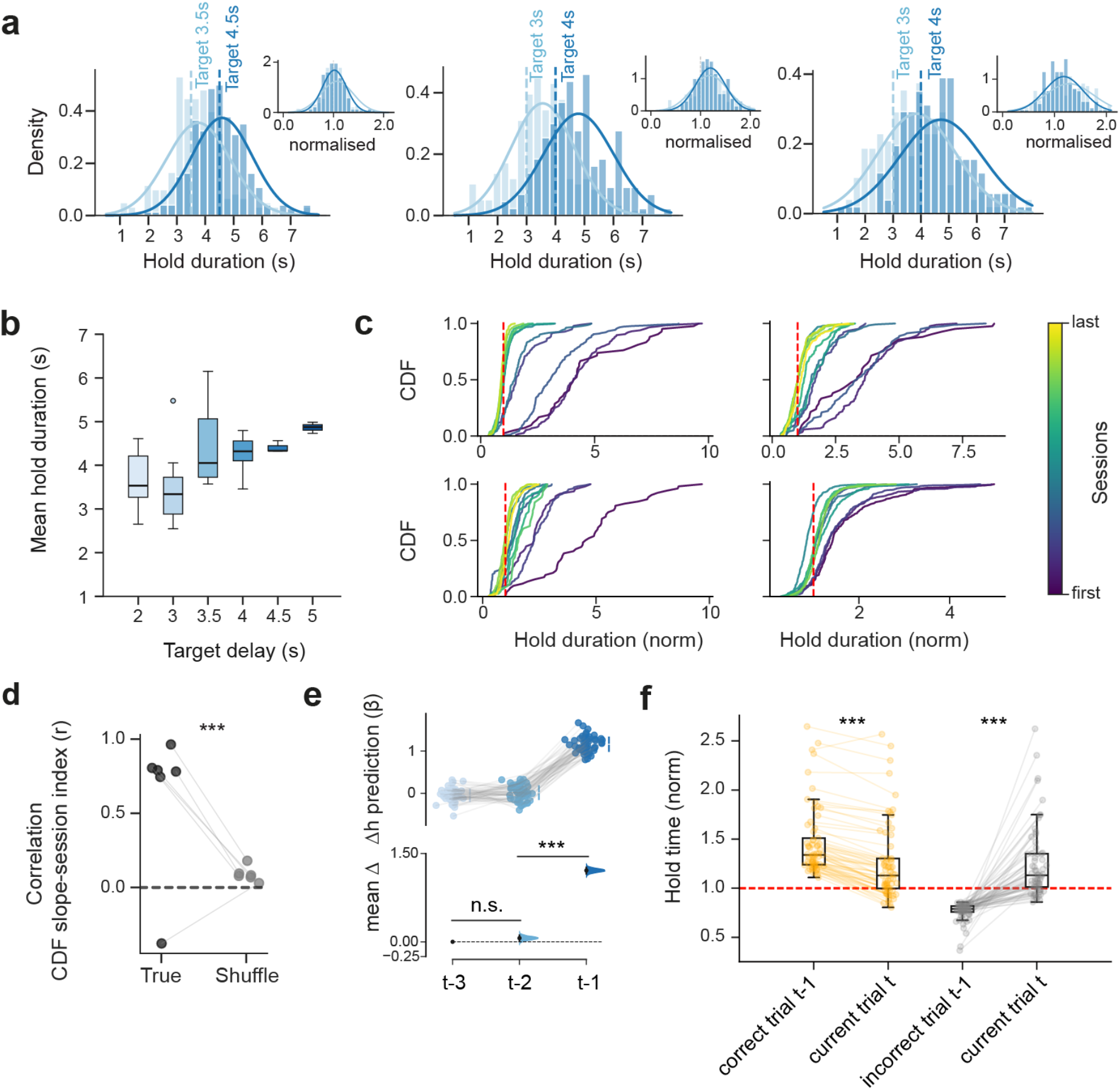
Behavioural adaptation and policy optimization across sessions. **(a)** Example distributions of bar-hold durations from three mice. Each panel shows two sessions (light and dark blue) with different target delays (dashed vertical lines). Insets show hold times normalized to the session target. **(b)** Mean bar-hold duration across sessions for each target delay (above 1 s). Boxes indicate median and interquartile range. **(c)** Cumulative distribution functions (CDFs) of normalized hold times from four example mice (one panel per mouse). Within each panel, sessions are color-coded by training order (i.e., session index; from early to late). CDFs become steeper around the target (dashed line; normalized time = 1) across sessions. **(d)** Mouse-wise Spearman correlation between session index and CDF slope. Across mice, CDFs became significantly steeper with training compared to surrogate data (see Methods; p = 7 × 10^-5^; one-tailed paired bootstrap test; n = 6 mice). **(e)** Regression coefficients from the trial-history linear regression model, corresponding to outcomes of the three preceding trials (*t* − 3, *t* − 2, *t* − 1). The most recent trial (t–1) carried more information than t–2 and t–3 (p < 10⁻⁵; paired bootstrap tests; Bonferroni corrected), which did not differ from each other (p = 0.07; paired bootstrap test; Bonferroni corrected; n = 56 sessions from 6 mice). **(f)** Change in normalized hold time (Δℎ; see Fig. 1e) on trial *t* as a function of outcome of the preceding trial *t* − 1 (correct, orange; incorrect, gray). Following correct trials, hold durations decreased; following incorrect trials, they increased (p < 10^-5^; paired bootstrap tests). Red dashed line indicates the target delay. *** p < 0.001. Data in b,f are from n = 65 sessions from 6 mice.

**Fig. S2:**
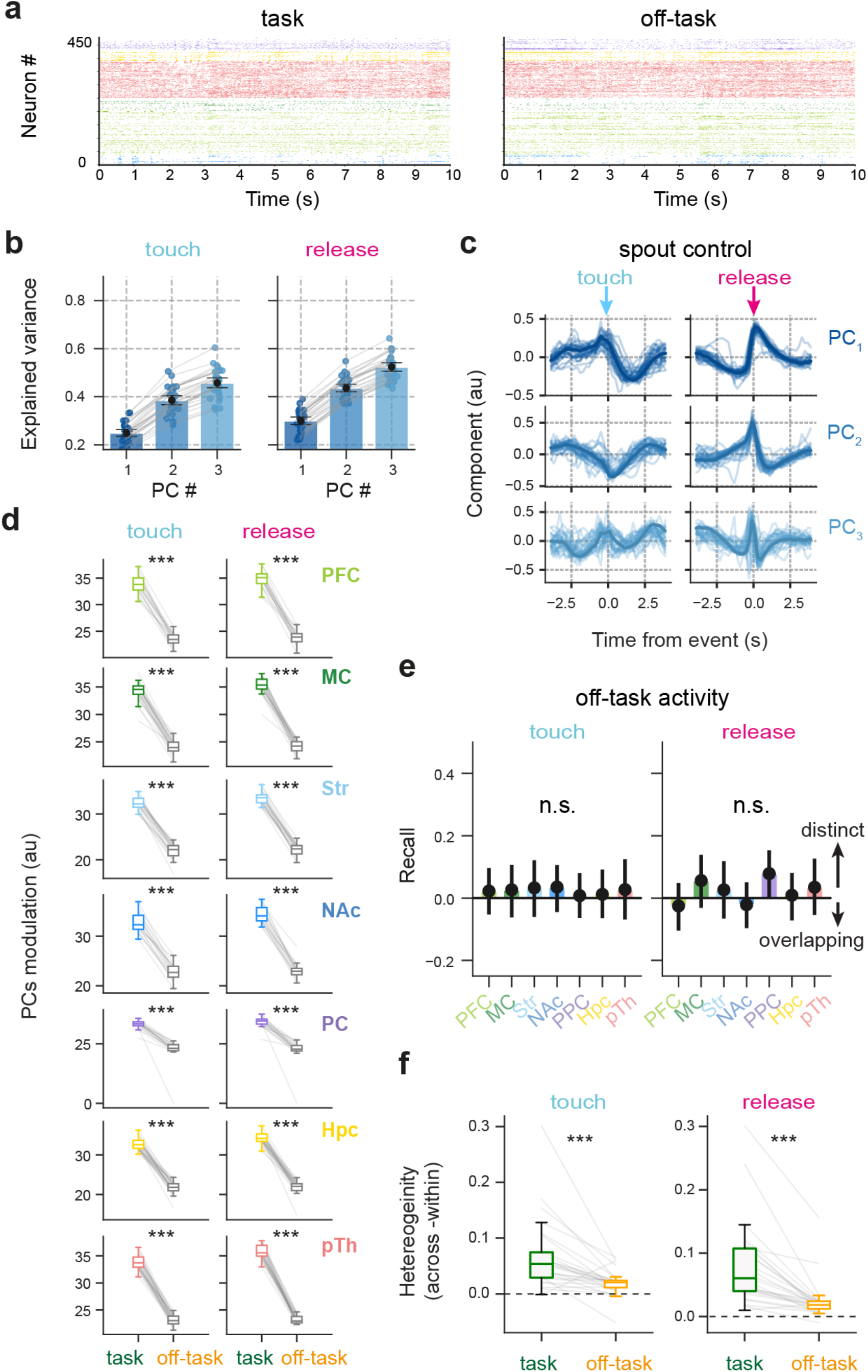
Principal component structure and regional heterogeneity of task-evoked responses. **(a)** Population rasters during 10-s epochs of task and off-task activity. Time is shown relative to epoch onset. **(b)** Variance explained by the first three principal components (PCs) of response waveforms (from Fig. 2b), aligned to handbar touch (left) and release (right). Each point represents one session. Cumulatively, the first three PCs explained more than 45% of the variance (mean variance explained [95% CI]: handbar touch, 46 [44–48] %; handbar release, 52 [51–54] %). Error bars, 95% CI. **(c)** First three PCs (one per row) for handbar touch (left) and release (right), computed using only trials not preceded by a correct trial to control for reward-related effects. Thin lines, individual sessions; thick lines, PCs extracted from the full dataset. **(d)** Modulation strength of response waveforms (see Methods) grouped by region (rows). Session-mean values are shown for task versus off-task control windows. All regions showed stronger modulation during task than off-task epochs (p < 10^-5^; paired bootstrap test; Bonferroni corrected). **(e)** Chance-normalised recall of an LDA classifier trained to discriminate region identity from PC-space representations derived from off-task windows (see Methods). Recall for each region did not differ from chance (recall > 0; p > 0.32; one-tailed bootstrap tests, Bonferroni corrected). Error bars, 95% CI across 1,000 permutations with equal sampling of neurons per region for classifier training. **(f)** Difference in response heterogeneity (across-region minus within-region) computed during task and off-task control windows. Heterogeneity was greater during task than off-task epochs (p < 10^-5^; paired bootstrap tests). * p < 0.05, ** p < 0.01, *** p < 0.001. Data in a–b,d are from n = 31 sessions from 6 mice; data in c–e are from n = 26 sessions from 5 mice.

**Fig. S3:**
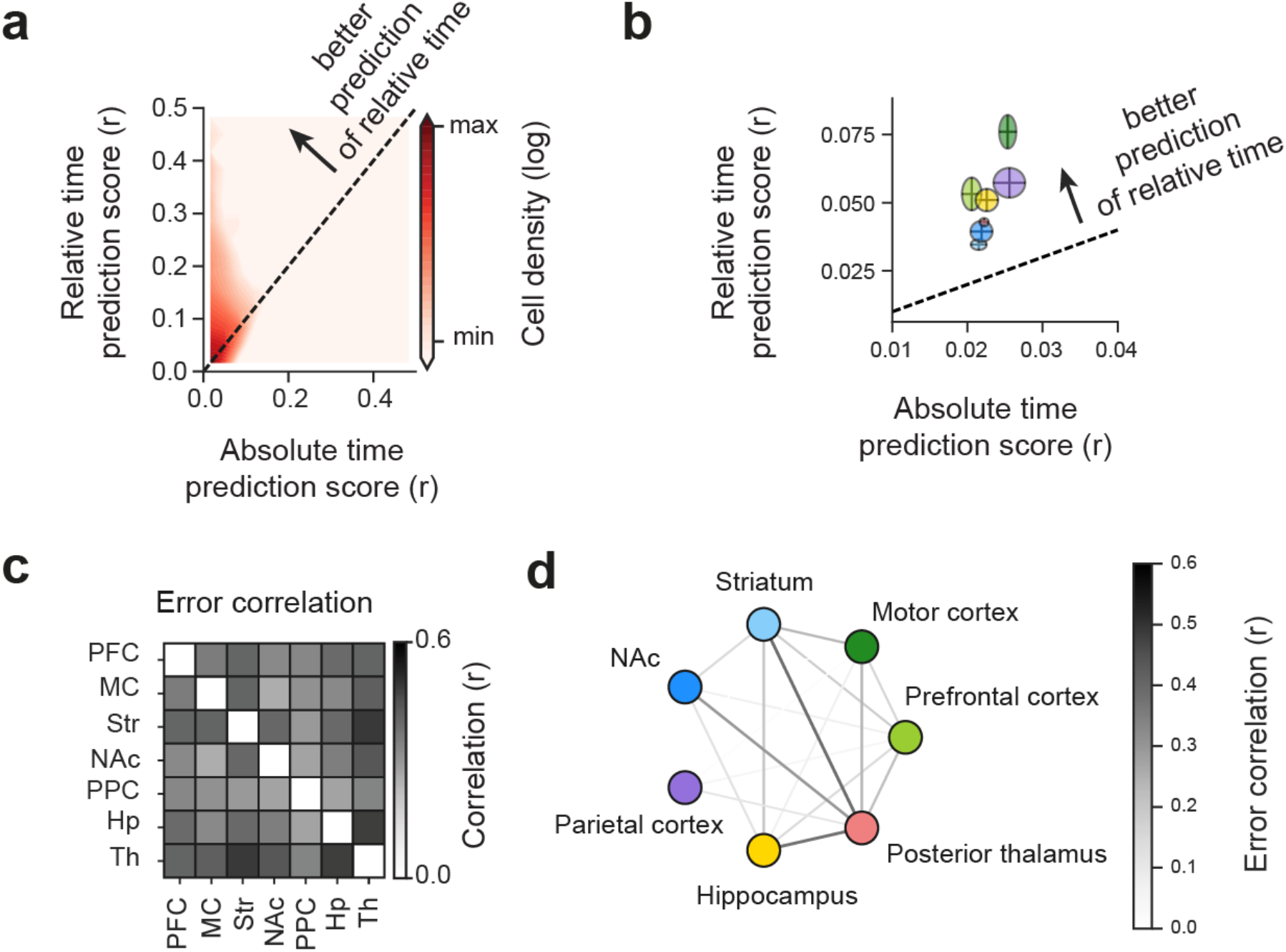
Single-neuron encoding of elapsed time and cross-regional decoding error correlations (a, **b)** Single-neuron encoding of absolute versus relative elapsed time. **(a)** Density heatmap of neurons plotted by decoding accuracy for absolute versus relative elapsed time (see Methods). Neurons above the diagonal preferentially encoded relative time, whereas those below preferentially encoded absolute time. Across the full dataset, the distribution was biased toward relative time encoding. **(b)** Same analysis shown by region. Ellipses indicate 95% CI for each region. All regions clustered above the diagonal, showing preferential encoding of relative time. **(c, d)** Regional error-correlation analysis. Trial-by-trial decoding errors for relative elapsed time were used to construct inter-regional error-correlation matrices. **(c)** Mean error-correlation matrix across sessions. **(d)** Graph representation of the matrix in (c), with nodes representing individual regions with edge thickness and gray scale indicating correlation magnitude. * p < 0.05, ** p < 0.01, *** p < 0.001. Data in a,b are from N = 8,642 neurons across the full dataset. Data in d,e are averaged across n = 31 sessions from 6 mice.

**Fig. S4:**
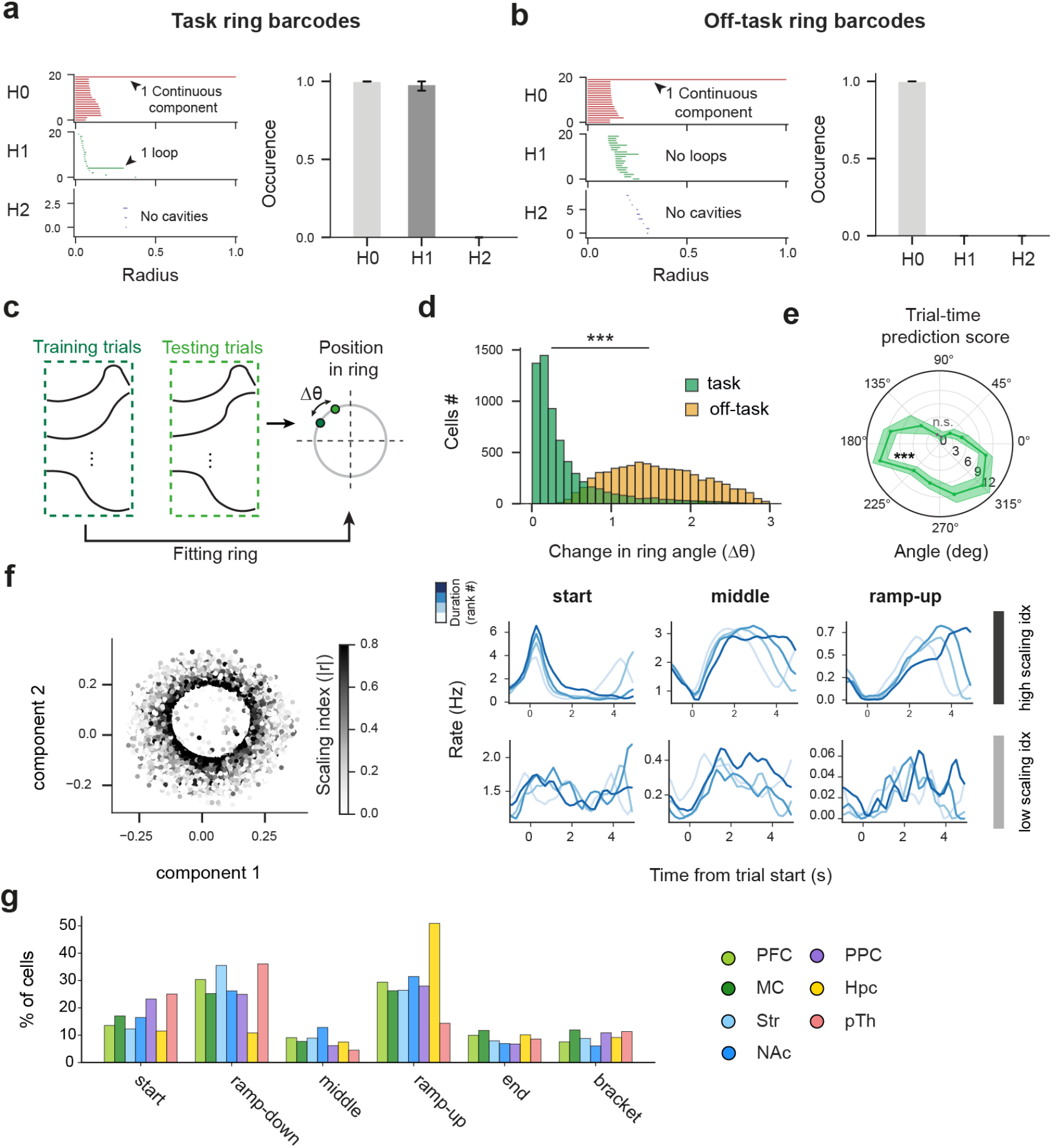
Topological validation and stability of the ring manifold. (a, **b)** Persistent homology analysis of the ring manifold. **(a)** Barcodes of the task-related manifold (left) showing one H0 and one H1 component, consistent with a 2D ring topology. The persistence of H0 and H1 components was maintained across 1,000 permutations of neurons within the manifold (right; p < 0.001 one-tailed bootstrap test; Bonferroni corrected). **(b)** Same analysis for manifolds constructed from off-task windows. The topology was consistent with a point cloud, showing only an H0 component (p < 0.001; bootstrap test; Bonferroni corrected). *Error bars*, 95% CI across 1,000 neuron permutations. **(c, d)** Stability of neuronal angular position within the ring manifold. **(c)** Schematic of stability analysis. Half of the trials were used to extract the warped-time tuning curve for each neuron, and their mean was used to fit the manifold and define training angular positions. Held-out trials were projected onto the ring to obtain test angular positions. The angular difference between training and testing (Δ*θ*) quantified stability. **(d)** Angular shift (Δ*θ*) for manifolds constructed from task versus off-task responses. Angular positions were more stable during task epochs (p < 0.001; paired bootstrap test). **(e)** Polar plot of chance-normalized relative-time decoding accuracy (framework from Fig. 3) for neurons grouped by ring angular sector. Neurons from most sectors decoded time above chance, except those between 45° and 105° (mainly associated with middle neurons) (p > 0.33; bootstrap test; n = 31 sessions from 6 mice). *Shaded areas*, 95% CI. **(f)** Ring manifold color-coded by scaling index (see Methods). Each dot represents one neuron (left). Right, example responses of six neurons grouped by high (top) or low (bottom) scaling index, spanning start, middle, and ramp-up profiles (columns). Traces show mean firing rate aligned to handbar touch for quartiles of trial duration (light to dark blue). **(g)** Mean proportion of neurons per region (color-coded) classified as start, ramp-down, middle, ramp-up/end, or bracket. All regions contributed to each profile, with regional biases (e.g., ramp-down enriched in posterior thalamus (pTh), ramp-up in hippocampus (Hpc), bracket in motor cortex (MC). * p < 0.05, ** p < 0.01, *** p < 0.001.

**Fig. S5:**
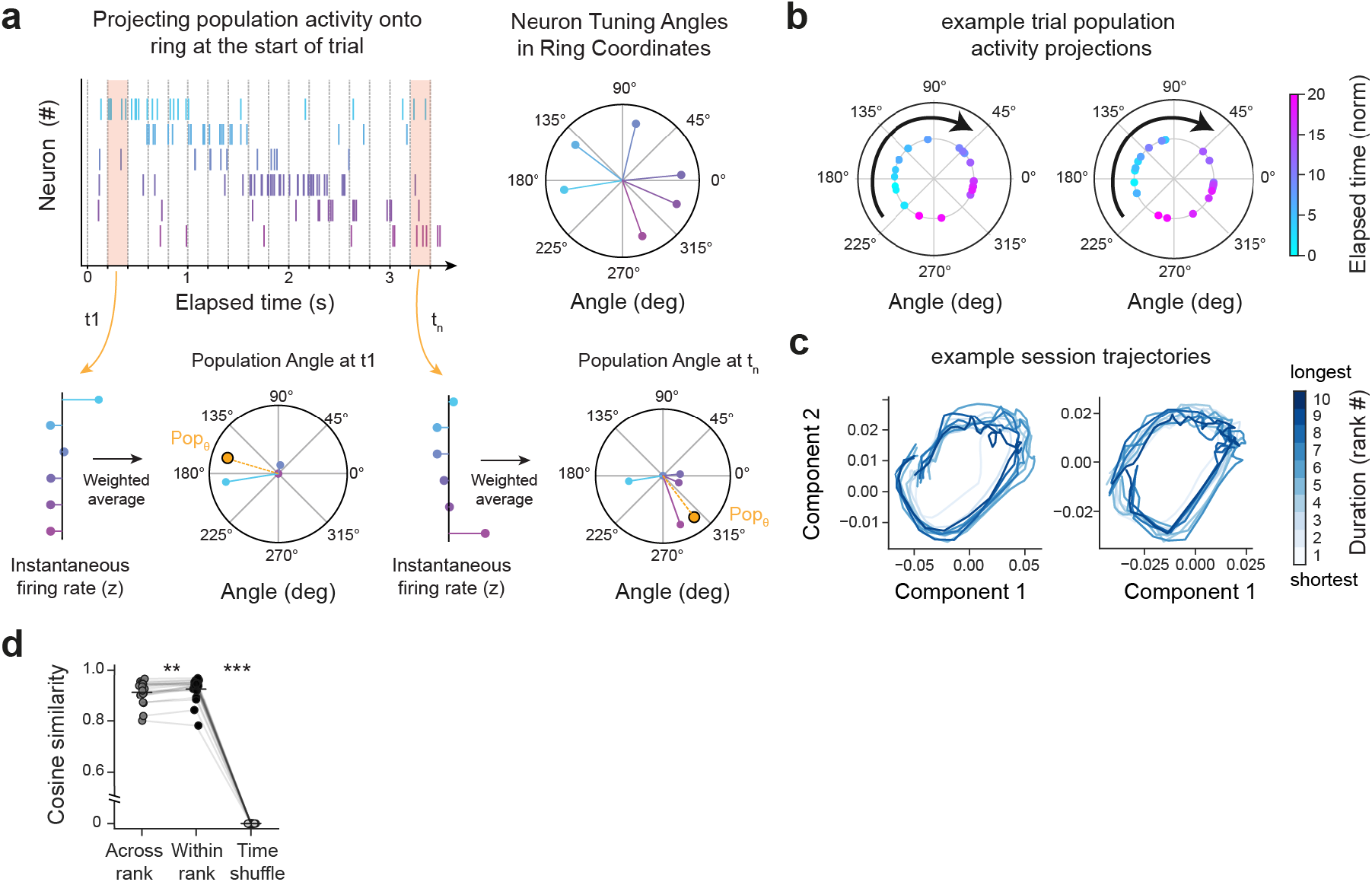
Angular population projection and trajectory reconstruction in ring space. **(a)** Validation of projection of instantaneous population activity onto ring-angular space (see Methods, “Elapsed-time prediction from low-dimensional angular trajectory”). Six simulated neurons with Gaussian-shaped warped-time tuning curves (left) were projected onto the ring manifold (from Fig. 4), distributing along the angular axis (right). Neurons tiled the warped interval uniformly from start to end (color-coded cyan to orchid). Simulated spiking activity for trials of different durations generated using Poisson firing based on the tuning curves in a. Instantaneous population activity (time-resolved firing rate vector) was used to compute a weighted mean of neuronal angular positions, yielding a population angular projection (*Pop_θ_*). The population angular projection evolved smoothly over time, forming an angular trajectory consistent with the simulated tuning structure. **(b)** Example single-trial angular trajectories from two different sessions. Each point represents a time bin within the trial, color-coded by elapsed time. **(c)** Mean angular trajectories from two example sessions, averaged across ten percentile bins of hold duration (shortest to longest). Note that trajectories overlapped in low-dimensional space despite differences in absolute trial duration. **(d)** Trajectory similarity within and across duration ranks. Cosine similarity between population trajectories was computed within duration ranks, across duration ranks, and after temporal shuffling of rank-averaged trajectories. Each point is a session; Similarity was high in both conditions, although higher within ranks, and both exceeded the shuffle control (paired Wilcoxon signed-rank tests, p < 0.001). * p < 0.05, ** p < 0.01, *** p < 0.001.

**Fig. S6:**
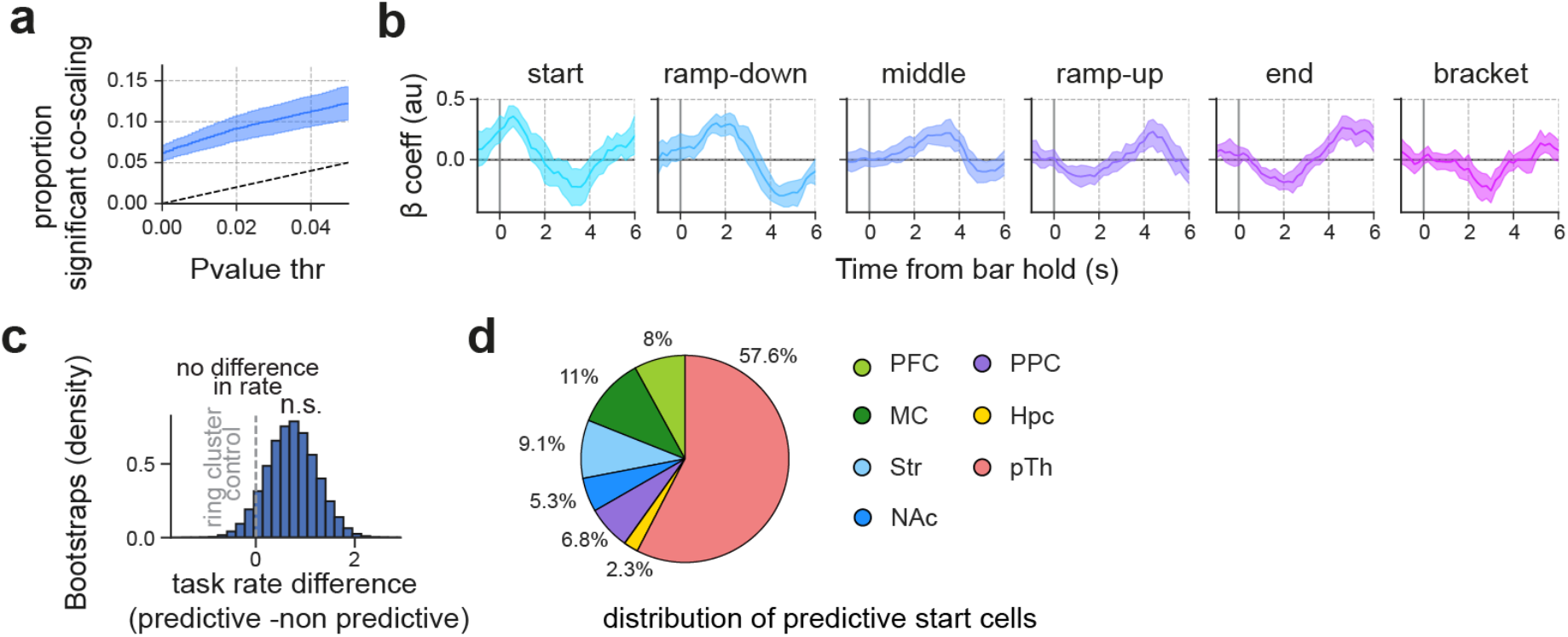
Characterization of trial-duration–predictive start neurons. **(a)** Proportion of neurons that significantly co-scaled their activity across trials with the rest of the population. The y-axis shows the observed proportion of neurons exceeding a given p-value threshold (x-axis), based on surrogate distributions controlling for independent scaling. The dashed diagonal indicates the expected false discovery rate. The observed proportion exceeds the diagonal, indicating more co-scaling neurons than expected by chance. Shaded area, 95% CI (n = 31 sessions from 5 mice). **(b)** Temporal evolution of regression coefficients for distinct ring profiles in predicting trial duration relative to handbar touch (see Methods). Coefficients different from zero indicate predictive information. The start ensemble showed elevated positive coefficients early in the trial (peaking within ∼1 s), consistent with early duration prediction. Shaded area, 95% CI (n = 13 sessions from 6 mice). **(c)** Bootstrapped mean difference in firing rate during the task between trial-duration– predictive and non-predictive start neurons (predictive minus non-predictive). No significant difference was observed (p = 0.35; bootstrap test; n = 264 predictive neurons, 257 non-predictive neurons), indicating comparable firing rates within the start cluster. **(d)** Distribution of trial-duration–predictive start neurons across regions (color-coded). The largest proportion was observed in posterior thalamus (pTH) and the smallest in hippocampus (Hpc); n = 297 predictive start neurons. * p < 0.05, ** p < 0.01, *** p < 0.001.

**Table. S1:**
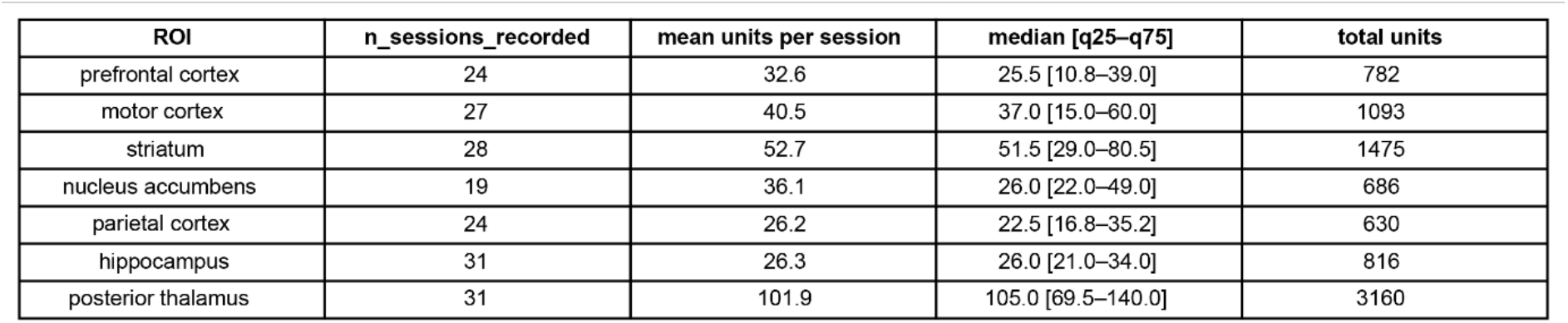
Distribution of recorded units across brain regions.

**Table. S2:** Parameters used for unit quality control with Bombcell. Thresholds and values applied for the classification and selection of single units across all sessions.

| Parameter | Value | Units |
| --- | --- | --- |
| waveformBaselineNoiseWindow | 20 | samples |
| tauR_valuesMin | 0.0015 | s |
| tauR_valuesStep | 0.0005 | s |
| tauR_valuesMax | 0.0015 | s |
| tauC | 0.0001 | s |
| hillOrLlobetMethod | 1 | — |
| computeTimeChunks | 1 | — |
| deltaTimeChunk | 360 | s |
| presenceRatioBinSize | 60 | s |
| driftBinSize | 60 | s |
| computeDrift | 0 | — |
| minThreshDetectPeaksTroughs | 0.1 | relative |
| normalizeSpDecay | 1 | — |
| minWidthFirstPeak | 4 | samples |
| minMainPeakToTroughRatio | 10 | — |
| minWidthMainTrough | 5 | samples |
| ephys_sample_rate | 30000 | Hz |
| nChannels | 384 | channels |
| nSyncChannels | 0 | channels |
| computeDistanceMetrics | 0 | — |
| nChannelsIsoDist | 4 | channels |
| splitGoodAndMua_NonSomatic | 0 | — |
| maxNPeaks | 2 | — |
| maxNTroughs | 1 | — |
| somatic | 1 | — |
| minWvDuration | 100 | μs |
| maxWvDuration | 1000 | μs |
| minSpatialDecaySlope | -0.005 | a.u./μm |
| maxWvBaselineFraction | 0.3 | fraction |
| firstPeakRatio | 3 | — |
| isoDmin | 20 | — |
| lratioMax | 0.1 | — |
| ssMin | NaN | — |
| minAmplitude | 20 | μV |
| maxRPVviolations | 0.2 | fraction |
| maxPercSpikesMissing | 20 | % |
| minNumSpikes | 200 | spikes |
| maxDrift | 100 | μm |
| minPresenceRatio | 0.6 | fraction |
| minSNR | 1 | — |

## Notes

### Competing Interest Statement

The authors have declared no competing interest.

## References

1. Merchant, H., Harrington, D. L. & Meck, W. H. Neural Basis of the Perception and Estimation of Time. Annu. Rev. Neurosci. 36, 313–336 (2013).

2. Paton, J. J. & Buonomano, D. V. The Neural Basis of Timing: Distributed Mechanisms for Diverse Functions. Neuron 98, 687–705 (2018).

3. Buhusi, C. V. & Meck, W. H. What makes us tick? Functional and neural mechanisms of interval timing. Nat Rev Neurosci 6, 755–765 (2005).

4. Parker, K. L., Chen, K.-H., Kingyon, J. R., Cavanagh, J. F. & Narayanan, N. S. D_1_ -Dependent 4 Hz Oscillations and Ramping Activity in Rodent Medial Frontal Cortex during Interval Timing. J. Neurosci. 34, 16774–16783 (2014).

5. Zhang, Q., Weber, M. A. & Narayanan, N. S. Medial prefrontal cortex and the temporal control of action. Int Rev Neurobiol 158, 421–441 (2021).

6. Leon, M. I. & Shadlen, M. N. Representation of Time by Neurons in the Posterior Parietal Cortex of the Macaque. Neuron 38, 317–327 (2003).

7. Gouvêa, T. S. et al. Striatal dynamics explain duration judgments. eLife 4, e11386 (2015).

8. MacDonald, C. J., Lepage, K. Q., Eden, U. T. & Eichenbaum, H. Hippocampal “Time Cells” Bridge the Gap in Memory for Discontiguous Events. Neuron 71, 737–749 (2011).

9. Lusk, N., Meck, W. H. & Yin, H. H. Mediodorsal Thalamus Contributes to the Timing of Instrumental Actions. J Neurosci 40, 6379–6388 (2020).

10. Kim, J., Ghim, J.-W., Lee, J. H. & Jung, M. W. Neural Correlates of Interval Timing in Rodent Prefrontal Cortex. J. Neurosci. 33, 13834–13847 (2013).

11. Emmons, E. B. et al. Rodent Medial Frontal Control of Temporal Processing in the Dorsomedial Striatum. J. Neurosci. 37, 8718–8733 (2017).

12. Xu, M., Zhang, S., Dan, Y. & Poo, M. Representation of interval timing by temporally scalable firing patterns in rat prefrontal cortex. Proc. Natl. Acad. Sci. U.S.A. 111, 480–485 (2014).

13. Tiganj, Z., Jung, M. W., Kim, J. & Howard, M. W. Sequential Firing Codes for Time in Rodent Medial Prefrontal Cortex. Cerebral Cortex 27, 5663–5671 (2017).

14. Zhou, S., Masmanidis, S. C. & Buonomano, D. V. Neural Sequences as an Optimal Dynamical Regime for the Readout of Time. Neuron 108, 651–658.e5 (2020).

15. Narayanan, N. S. Ramping activity is a cortical mechanism of temporal control of action. Current Opinion in Behavioral Sciences 8, 226–230 (2016).

16. Janssen, P. & Shadlen, M. N. A representation of the hazard rate of elapsed time in macaque area LIP. Nat Neurosci 8, 234–241 (2005).

17. Cook, J. R. et al. Secondary auditory cortex mediates a sensorimotor mechanism for action timing. Nat Neurosci 25, 330–344 (2022).

18. Heys, J. G. & Dombeck, D. A. Evidence for a subcircuit in medial entorhinal cortex representing elapsed time during immobility. Nat Neurosci 21, 1574–1582 (2018).

19. Churchland, M. M. et al. Neural population dynamics during reaching. Nature 487, 51–56 (2012).

20. Mante, V., Sussillo, D., Shenoy, K. V. & Newsome, W. T. Context-dependent computation by recurrent dynamics in prefrontal cortex. Nature 503, 78–84 (2013).

21. Chaudhuri, R., Gerçek, B., Pandey, B., Peyrache, A. & Fiete, I. The intrinsic attractor manifold and population dynamics of a canonical cognitive circuit across waking and sleep. Nat Neurosci 22, 1512–1520 (2019).

22. Gallego, J. A. et al. Cortical population activity within a preserved neural manifold underlies multiple motor behaviors. Nat Commun 9, 4233 (2018).

23. Sadtler, P. T. et al. Neural constraints on learning. Nature 512, 423–426 (2014).

24. Nieh, E. H. et al. Geometry of abstract learned knowledge in the hippocampus. Nature 595, 80–84 (2021).

25. Gardner, R. J. et al. Toroidal topology of population activity in grid cells. Nature 602, 123–128 (2022).

26. Wang, J., Narain, D., Hosseini, E. A. & Jazayeri, M. Flexible timing by temporal scaling of cortical responses. Nat Neurosci 21, 102–110 (2018).

27. Vyas, S., Golub, M. D., Sussillo, D. & Shenoy, K. V. Computation Through Neural Population Dynamics. Annu. Rev. Neurosci. 43, 249–275 (2020).

28. Remington, E. D., Narain, D., Hosseini, E. A. & Jazayeri, M. Flexible Sensorimotor Computations through Rapid Reconfiguration of Cortical Dynamics. Neuron 98, 1005–1019.e5 (2018).

29. Mita, A., Mushiake, H., Shima, K., Matsuzaka, Y. & Tanji, J. Interval time coding by neurons in the presupplementary and supplementary motor areas. Nat Neurosci 12, 502– 507 (2009).

30. Matell, M. S. & Meck, W. H. Cortico-striatal circuits and interval timing: coincidence detection of oscillatory processes. Cognitive Brain Research 21, 139–170 (2004).

31. Soares, S., Atallah, B. V. & Paton, J. J. Midbrain dopamine neurons control judgment of time. Science 354, 1273–1277 (2016).

32. Schall, T. A. et al. Temporal dynamics of nucleus accumbens neurons in male mice during reward seeking. Nat Commun 15, 9285 (2024).

33. Heldman, R., Pang, D., Zhao, X., Mensh, B. & Wang, Y. Time or distance encoding by hippocampal neurons via heterogeneous ramping rates. Nat Commun 16, 11083 (2025).

34. Shimbo, A., Izawa, E.-I. & Fujisawa, S. Scalable representation of time in the hippocampus. Sci. Adv. 7, eabd7013 (2021).

35. Sherman, S. M. Thalamus plays a central role in ongoing cortical functioning. Nat Neurosci 19, 533–541 (2016).

36. Halassa, M. M. & Kastner, S. Thalamic functions in distributed cognitive control. Nat Neurosci 20, 1669–1679 (2017).

37. De Corte, B. J., Akdoğan, B. & Balsam, P. D. Temporal scaling and computing time in neural circuits: Should we stop watching the clock and look for its gears? Front. Behav. Neurosci. 16, 1022713 (2022).

38. Macar, F., Vidal, F. & Casini, L. The supplementary motor area in motor and sensory timing: evidence from slow brain potential changes. Exp Brain Res 125, 271–280 (1999).

39. Sebastian, E. R., Esparza, J. & M de la Prida, L. Quantifying the distribution of feature values over data represented in arbitrary dimensional spaces. PLoS Comput Biol 20, e1011768 (2024).

40. Mello, G. B. M., Soares, S. & Paton, J. J. A Scalable Population Code for Time in the Striatum. Current Biology 25, 1113–1122 (2015).

41. Sawatani, F., Ide, K. & Takahashi, S. The neural representation of time distributed across multiple brain regions differs between implicit and explicit time demands. Neurobiology of Learning and Memory 199, 107731 (2023).

42. Salz, D. M. et al. Time Cells in Hippocampal Area CA3. Journal of Neuroscience 36, 7476–7484 (2016).

43. MacDonald, C. J., Carrow, S., Place, R. & Eichenbaum, H. Distinct Hippocampal Time Cell Sequences Represent Odor Memories in Immobilized Rats. J. Neurosci. 33, 14607–14616 (2013).

44. Pastalkova, E., Itskov, V., Amarasingham, A. & Buzsáki, G. Internally generated cell assembly sequences in the rat hippocampus. Science 321, 1322–1327 (2008).

45. Gill, P. R., Mizumori, S. J. Y. & Smith, D. M. Hippocampal episode fields develop with learning. Hippocampus 21, 1240–1249 (2011).

46. Yuan, L. et al. Time cell sequences during delay intervals are not dependent on brain state and do not support hippocampus-dependent working memory. Nat Commun 16, 7470 (2025).

47. Jog, M. S., Kubota, Y., Connolly, C. I., Hillegaart, V. & Graybiel, A. M. Building Neural Representations of Habits. Science 286, 1745–1749 (1999).

48. Cunningham, P. J., Regier, P. S. & Redish, A. D. Dorsolateral Striatal Task-initiation Bursts Represent Past Experiences More than Future Action Plans. J Neurosci 41, 8051– 8064 (2021).

49. Buonomano, D. V. & Laje, R. Population clocks: motor timing with neural dynamics. Trends Cogn Sci 14, 520–527 (2010).

50. Ivry, R. B. & Schlerf, J. E. Dedicated and intrinsic models of time perception. Trends Cogn Sci 12, 273–280 (2008).

51. Tsao, A., Yousefzadeh, S. A., Meck, W. H., Moser, M.-B. & Moser, E. I. The neural bases for timing of durations. Nat Rev Neurosci 23, 646–665 (2022).

52. Harvey, C. D., Coen, P. & Tank, D. W. Choice-specific sequences in parietal cortex during a virtual-navigation decision task. Nature 484, 62–68 (2012).

53. Tsao, A., Yousefzadeh, S. A., Meck, W. H., Moser, M.-B. & Moser, E. I. The neural bases for timing of durations. Nat Rev Neurosci 23, 646–665 (2022).

54. Remington, E. D., Narain, D., Hosseini, E. A. & Jazayeri, M. Flexible Sensorimotor Computations through Rapid Reconfiguration of Cortical Dynamics. Neuron 98, 1005–1019.e5 (2018).

55. Maimon, G. & Assad, J. A. A cognitive signal for the proactive timing of action in macaque LIP. Nat Neurosci 9, 948–955 (2006).

56. Murakami, M., Vicente, M. I., Costa, G. M. & Mainen, Z. F. Neural antecedents of self-initiated actions in secondary motor cortex. Nat Neurosci 17, 1574–1582 (2014).

57. Buonomano, D. V. & Laje, R. Population clocks: motor timing with neural dynamics. Trends Cogn Sci 14, 520–527 (2010).

58. Jones, C. R. G. & Jahanshahi, M. Dopamine Modulates Striato-Frontal Functioning during Temporal Processing. Front. Integr. Neurosci. 5, (2011).

59. Meck, W. H. Neuropharmacology of timing and time perception. Cognitive Brain Research 3, 227–242 (1996).

60. Agostino, P. V. & Cheng, R.-K. Contributions of dopaminergic signaling to timing accuracy and precision. Current Opinion in Behavioral Sciences 8, 153–160 (2016).

61. Maricq, A. V., Roberts, S. & Church, R. M. Methamphetamine and time estimation. Journal of Experimental Psychology: Animal Behavior Processes 7, 18–30 (1981).

62. Otchy, T. M. et al. Acute off-target effects of neural circuit manipulations. Nature 528, 358–363 (2015).

63. Wolff, S. B. & Ölveczky, B. P. The promise and perils of causal circuit manipulations. Current Opinion in Neurobiology 49, 84–94 (2018).

64. Guo, Z. V. et al. Maintenance of persistent activity in a frontal thalamocortical loop. Nature 545, 181–186 (2017).

65. Inagaki, H. K., Fontolan, L., Romani, S. & Svoboda, K. Discrete attractor dynamics underlies persistent activity in the frontal cortex. Nature 566, 212–217 (2019).

66. Li, N., Daie, K., Svoboda, K. & Druckmann, S. Robust neuronal dynamics in premotor cortex during motor planning. Nature 532, 459–464 (2016).

67. Galiñanes, G. L., Bonardi, C. & Huber, D. Directional Reaching for Water as a Cortex-Dependent Behavioral Framework for Mice. Cell Reports 22, 2767–2783 (2018).

68. Peng, Y. et al. Brain-wide population activity during reaching integrates action-mediated goal expectation. Preprint at 10.1101/2024.11.04.621878 (2024).

69. The International Brain Laboratory et al. Standardized and reproducible measurement of decision-making in mice. eLife 10, e63711 (2021).

70. Akam, T. et al. Open-source, Python-based, hardware and software for controlling behavioural neuroscience experiments. eLife 11, e67846 (2022).

71. Leon, M. I. & Shadlen, M. N. Representation of Time by Neurons in the Posterior Parietal Cortex of the Macaque. Neuron 38, 317–327 (2003).

72. Narayanan, N. S. & Laubach, M. Delay Activity in Rodent Frontal Cortex During a Simple Reaction Time Task. Journal of Neurophysiology 101, 2859–2871 (2009).

73. Shamash, P., Carandini, M., Harris, K. D. & Steinmetz, N. A. A tool for analyzing electrode tracks from slice histology. Preprint at 10.1101/447995 (2018).

74. Liu, L. D. et al. Accurate Localization of Linear Probe Electrode Arrays across Multiple Brains. eNeuro 8, ENEURO.0241-21.2021 (2021).

75. Pachitariu, M., Sridhar, S. & Stringer, C. Solving the spike sorting problem with Kilosort. Preprint at 10.1101/2023.01.07.523036 (2023).

76. Jake Pennington et al. MouseLand/Kilosort: Kilosort v4.1.3. Zenodo 10.5281/ZENODO.3597474 (2025).

77. Fabre, J. M. J., Van Beest, E. H., Peters, A. J., Carandini, M. & Harris, K. D. Bombcell: automated curation and cell classification of spike-sorted electrophysiology data. Zenodo 10.5281/ZENODO.8172821 (2023).

78. Harris, C. R. et al. Array programming with NumPy. Nature 585, 357–362 (2020).

79. McKinney, W. & others. Data structures for statistical computing in Python. Scipy 445, 51–56 (2010).

80. Hunter, J. D. Matplotlib: A 2D graphics environment. Computing in science & engineering 9, 90–95 (2007).

81. Waskom, M. seaborn: statistical data visualization. JOSS 6, 3021 (2021).

82. Virtanen, P. et al. SciPy 1.0: fundamental algorithms for scientific computing in Python. Nat Methods 17, 261–272 (2020).

83. Pedregosa, F. et al. Scikit-learn: Machine learning in Python. the Journal of machine Learning research 12, 2825–2830 (2011).

84. Hagberg, A. A., Schult, D. A. & Swart, P. J. Exploring Network Structure, Dynamics, and Function using NetworkX. in 11–15 (Pasadena, California, 2008). doi:10.25080/TCWV9851.

85. Seabold, S. & Perktold, J. Statsmodels: Econometric and Statistical Modeling with Python. in 92–96 (Austin, Texas, 2010). doi:10.25080/Majora-92bf1922-011.

86. Ho, J., Tumkaya, T., Aryal, S., Choi, H. & Claridge-Chang, A. Moving beyond P values: data analysis with estimation graphics. Nat Methods 16, 565–566 (2019).

